# Electroencephalogram (EEG) Classification using a bio-inspired Deep Oscillatory Neural Network

**DOI:** 10.1101/2024.05.24.595714

**Authors:** Sayan Ghosh, C. Vigneswaran, NR Rohan, V.Srinivasa Chakravarthy

**Affiliations:** Indian Institute of Technology Madras

**Keywords:** Deep Oscillatory Neural Network, Oscillatory CNN, EEG signal, Hopf Oscillator, Classification

## Abstract

In this paper, we propose two models of oscillatory neural networks - the Deep Oscillatory Neural Network (DONN) and a convolutional variation of it named Oscillatory Convolutional Neural Network (OCNN) – and apply the models to a variety of problems involving the classification and prediction of Electroencephalogram (EEG) signals. Deep neural networks applied to signal processing problems will have to incorporate various architectural features to remember the history of the input signals e.g., loops between the layers, “gated” neurons, and tapped delay lines. But real brains have rich dynamics expressed in terms of frequency bands like alpha, beta, gamma, delta, etc. To incorporate this aspect of brain dynamics in a Recurrent Neural Network (RNN) we propose to use nonlinear oscillators as dynamic neuron models in the hidden layers. The two oscillatory deep neural networks proposed are applied to the following EEG classification and prediction problems: Prediction of nearby EEG channels, classification of single-channel EEG data (healthy vs. epileptic, different stages of sleep stage classification), and multi-channel EEG data (Epileptic vs. Normal, Left vs. right-hand Motor imagery movement, and healthy vs. Claustrophobic EEG).

## 1. Introduction

Recent advancements in computational facilities and machine learning tools have shown a great impact on biosignal analysis with healthcare applications. One such example is the development in the field of electroencephalogram (EEG) technology and its diverse applications in brain-computer interface [10], detection of diseases [3, 33, 39], emotion recognition [5,7], etc. A majority of instances where machine learning tools have been applied to EEG processing involve some kind of signal classification. There is nearly a three-decade-long history of application of neural networks for EEG classification [39, 46-47]. Developments in neural network theory at the architecture level and single neuron level have contributed to the steady progress in this area.

Since EEG classification is a sequence processing problem, the concerned classifier must capture the history of the signal in some form. This can be done either at the level of the signal itself by inserting a tapped delay line at the input, or by adding memory mechanisms in the network. There were several instances of time-delay neural networks applied to EEG classification [1-2,6]. Early attempts to create neural networks with memory used various kinds of loops, resulting in the earliest Recurrent Neural Networks (RNNs), which were extensively applied to EEG processing [49-50]. However, RNNs in their original form suffered from the problem of vanishing gradients, which posed challenges in sequence processing with long time horizons [53].

Development of a special class of neuron models that may be collectively described as “gating” neuron models - Long Short Term Memory neurons (LSTM) [4], Gated Recurrent Unit (GRU) [4], Flip-flop neuron [48,76] – has immensely enhanced the performance of RNNs [4]. LSTMs and GRUs have shown greater performance over traditional RNNs while classifying sets of emotion from EEG signals [4]. There are many other applications of LSTM-type networks such as cognitive load prediction [5], emotion detection in Alzheimer’s disease (AD) conditions [7], epileptic seizure detection [2, 6], and motor imagery studies [10,29-30]. An RNN-BiLSTM network was used to classify the UCI epileptic seizure data [6]. Spatio-temporal feature fusion using CNN-LSTM has been applied to the problem of classifying motor imagery using BCI 2a dataset [10,35]. Hybrid RNN-LSTM was applied to the diagnosis of schizophrenia using EEG signals [3].

In this work, we apply a Deep Oscillatory Neural Network (DONN) to a variety of problems involving EEG time series prediction and classification. The DONN model has been recently formulated by our group as a generalization over the traditional deep neural networks with sigmoidal neurons [55]. In current RNN models, the most popular example of a single neuron/unit model exhibiting dynamics is a “gating” neuron [52]. RNNs with gating neurons exhibit excellent sequence processing capability but are rather weak in terms of biological plausibility. Neural activity in the real brain exhibits rich dynamics that may be resolved in terms of various frequency bands like alpha, beta, gamma, theta, etc [51]. No such oscillations may be discovered in RNNs with LSTM. Intending to develop networks that possess sequence processing capabilities and can explicitly exhibit oscillations, we developed the DONNs in which single neurons are modeled by nonlinear oscillators.

Also one major drawback of the RNN-type network is error propagation through cascaded network [77] and computational cost during training with this motivation we presently apply DONNs for various EEG classifications as well as prediction problems and demonstrate results comparable to the state-of-the-art performance by RNNs. To the best of our knowledge, this is the 1^st^ study where hopf oscillatory network is used to classify different sets of EEG data. We present two variations of the DONN. In the first, all the layers are 1D layers; but the layers are hybrid layers: some layers have oscillatory neurons while some have traditional sigmoidal neurons. The other model is the Oscillatory Convolutional Neural Network (OCNN), fashioned after the Convolutional Neural Network. We also show that in the OCNN model, the feature maps act as spatiotemporal filters that can process time-varying image data i.e. video data. Thus, we apply OCNN to classify topographical map sequences of EEG. We show in this study our DONN and OCNN models have enormous potential in the field of EEG technology with applications ranging from clinical to bio-robotics domains.

The aforementioned models of oscillatory networks (DONN and OCNN) are applied to the following problems in EEG processing:

1. Epilepsy EEG data (IIT D Epilepsy EEG data, BONN Dataset, CHB-MIT Dataset)
2. Motor Imagery EEG data
3. Sleep EEG (Supplementary file)
4. Claustrophobic EEG (Supplementary file)

The outline of the paper is as follows. Section 2 presents detailed model specifications and the data sets used. Section 3 describes the results which are discussed in the final section.

## 2. Methods

### 2.1 Data sets

We use the following EEG data sets – A, B, C, D, E (section 2.1.1-2.1.5) – for our classification studies. Data sets A-B (2.1.1-2.1.2) are single channel epilepsy datasets and DONN models are used to classify them. In case of data sets C-D (2.1.3-2.1.4), which pertain to multi-channel epilepsy and motor imagery, we represent the data in an image format using the topoplot technique [20] and use a OCNN to classify data. We also show that the DONN model is capable of solving regression problems. To this end, we train the DONN on the dataset E (2.1.5) which a BCI-competition dataset, where we take 8 neighboring channels as inputs and predict the channel located in the center of the 8 input channels.

#### 2.1.1 A. IITD Epilepsy Dataset

The IITD epilepsy data are recorded using Grass Telefactor Comet AS40 Amplification System from 10 epilepsy patients from Neurology & Sleep Centre, Hauz Khas, New Delhi [14, 73]. The dataset is recorded using a Comet AS40 EEG machine with a 200 Hz sampling rate for a duration of 5.12 Seconds. The dataset is segmented into preictal, interictal, and ictal, and filtered in the frequency range of 0.5 to 70 Hz, each set having 50 chunks of 1024 samples. For our study, we explored both two-class (pre-ictal vs. ictal EEG) and three-class classification problems (pre-ictal vs. inter-ictal vs. ictal). The dataset has 575 epileptic seizure segments and 529 non-epileptic seizure segments.

#### 2.1.2 B. BONN epilepsy Dataset

This dataset was recorded from the Department of Epileptology at the University of Bonn, Germany [15]. There are 5 sets of EEG data (A-E), containing 100 single channel EEGs of duration 23.6 sec, recorded at 173.61 Hz sampling frequency. The recorded EEG data are preprocessed by bandpass filtering (0.53 to 70 Hz). There are 4097 samples in each dataset. Sets A and B were collected on five healthy volunteers, who were relaxed in an awake state with eyes open (A) and eyes closed (B), respectively. From five patients, sets C and D were recorded during the seizure-free interval, and set E was recorded during seizure occurrence. In this present study, we have done six different combinations of classification studies.

B.1: Set A vs E: The combination of A and E datasets, here only A (relaxed, eye-open, non-ictal interval) and E (ictal interval) are used.

B.2: Set B vs E: The combination of B and E datasets, here only B (relaxed, eye closed, non-ictal interval) and E (ictal interval) are used.

B.3: Set C vs E: combination of C and E datasets, here only C (seizure-free interval) and E (ictal interval) are used.

B.4: Set D vs E: combination of D and E datasets, here only A (seizure-free interval) and E (ictal interval) are used.

B.5: Set C vs D: combination of C and D datasets.

B.6: Set A vs C vs E: combination of A, C, E datasets.

#### 2.1.3 C. CHB MIT Dataset

CHB-MIT (Children’s Hospital Boston) dataset is a repository of intractable seizure EEG data of 23 paediatric patients with an age group of 1.5 to 22 years including 5 male and 17 female subjects [17, 75]. The dataset is recorded with 256 Hz sampling frequency with 16-bit resolution and consists of 198 seizure events annotated by epilepsy experts. This dataset consists of 916-hour-long EEG data distributed over 664 EEG files. The EEG signals were segmented by the timing window since the data are long-hour data. Most of the files contained 23-channel EEG recordings and a few of the records had 24-26 channel recordings. The recording system follows the standard 10-20 electrode placement rule. The dataset contains bipolar EEG electrodes (FP1-F7, F7-T7, T7-P7, P7-O1, FP1-F3, F3-C3, C3-P3, P3-O1, FP2-F4, F4-C4, C4-P4, P4-O2, FP2-F8, F8-T8, P8-O2, FZ-CZ, CZ-PZ, P7-T7, T7-FT9, FT9-FT10, FT10-T8, and T8-P8) [18]. To standardize the model, we used 19 channels excluding P7-O1, P3-O1, T7-FT9, and T8-P8. There is a total of 6083 sec of seizure activity from all the subjects. The signal is pre-processed with a bandpass filter (between 0.1 to 70 Hz) and a notch filter (at 50 Hz) to remove the line noise.

#### 2.1.4 D. Motor Imagery EEG Dataset for classification

Motor imagery EEG data are recorded from 13 individuals (8 male and 5 female) from Toros University and Mersin University in the city of Mersin, Turkey [19]. The EEG data are recorded at 200 Hz,19 channels (‘Fp1’, ‘Fp2’, ‘F3’, ‘F4’, ‘C3’, ‘C4’, ‘P3’, ‘P4’, ‘O1’, ‘O2’, ‘F7’, ‘F8’, ‘T3’, ‘T4’, ‘T5’, ‘T6’, ‘Fz’, ‘Cz’, ‘Pz’) EEG-1200 EEG system. The participants have to implement mental imagery based on visual stimuli presented on the computer screen. One motor imagery experiment took approximately 1sec duration. Three different motor imagery tasks are performed here: Classical left/right motor imagery hand movement (CLA), HaLT (left, right leg, and tongue), and NoMT (movement of five fingers in one hand). In this study, we used CLA datasets. Each stimulus was shown for 1 second during which, participants had to perform a corresponding motor imagery task. EEG data are filtered in between [8 to 30 Hz].

#### 2.1.5 E. BCI Dataset -2A

We used a publicly available BCI competition IV2a [13] dataset for the neighbouring EEG channel prediction task. This task consists of using a set of 8 channels from the periphery of a 3X3 array of electrode locations and predicting the central channel. This problem is selected since it is an interesting regression problem compared to all the previous ones which have been classification problems. The aim is to demonstrate that the DONN model can perform both classification and regression tasks. This dataset has 22 EEG channels. Three motor imagery tasks (left hand, right hand, tongue) have been performed here. 4 healthy participants were included in this study. The EEG signals were sampled at 250 Hz, and filtered by a bandpass filter of 0.5 to 10 Hz.

## 3. Architecture of the Deep Oscillatory Neural Network (DONN)

The DONN model is essentially a feedforward network in which some layers have oscillatory neurons and other layers have traditional neurons like sigmoidal neurons, tanh neurons, ReLU neurons etc. Successive layers are connected by weight stages where the weights are complex-valued. Although the architecture is feedforward, the network is capable of sequence or signal processing since the oscillatory neurons have dynamics and therefore memory.

### 3.1 Oscillatory neuron model

The oscillatory neuron model we use in our networks is the Hopf oscillator [31]. It is a harmonic oscillator with a stable limit cycle. The canonical Hopf oscillator is described by the following complex-valued differential equation,

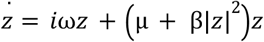

where *z* is the state variable given by

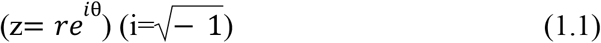

The polar coordinate representation of the above oscillator equation is:

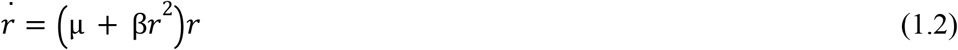

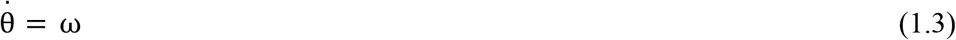

where *r* and θ are the state variables of the oscillator, 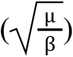 is the amplitude of oscillation, μ & β are bifurcation parameters and ω is the natural frequency of the oscillator.

I(t) is the external input signal, and ω is the angular frequency of the externally applied signal.

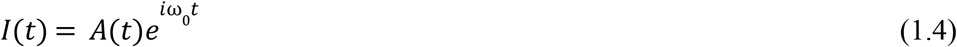

where Input presented as an additive input to the oscillators is shown below:

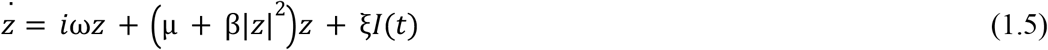

The polar coordinate representation of the last equation is:

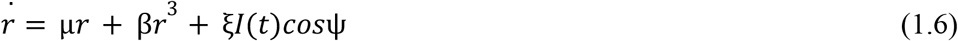

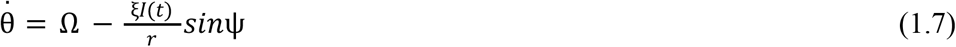

where Ψ = φ − ω_0_ *t* and Ω = ω − ω_0_ is the difference between the angular frequencies of the oscillators and the external input.

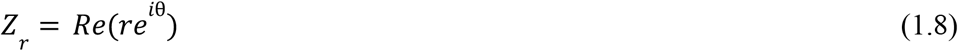

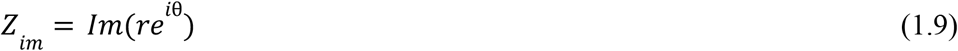

where *Z*_*r*_ and *Z*_*in*_ are the real and imaginary part of the oscillator’s state variable in complex domain.

We call this mode the resonator mode because when the input is a complex sinusoid, the oscillator responds with the highest amplitude when the input frequency matches the intrinsic frequency of the oscillator [54].

The oscillator shows resonance over a small range of frequencies around its natural frequency. The oscillator activation is then in a frequency locked state with the input signal i.e., the frequency of the oscillator output is equal to the frequency of the input signal. Furthermore, within a smaller range of frequency, around the resonant frequency, the oscillator activation is phase locked to the input signal. Beyond this regime, the system undergoes phase slip. However, the phase difference between the input signal and the intrinsic oscillations is bounded.

### 3.2 Network architecture

The architecture of the DONN model is shown in (fig. 2). This kind of general feedforward network model can be applied to regression as well as classification problems. In a typical EEG classification task, the input to the network is one or several EEG signals, and the output encodes the class identity. Fig.2 shows only a single input signal for simplicity; We have all-to-all connectivity from the input to the H1 layer (first hidden layer), which is a ReLU layer.

**Figure 1:**
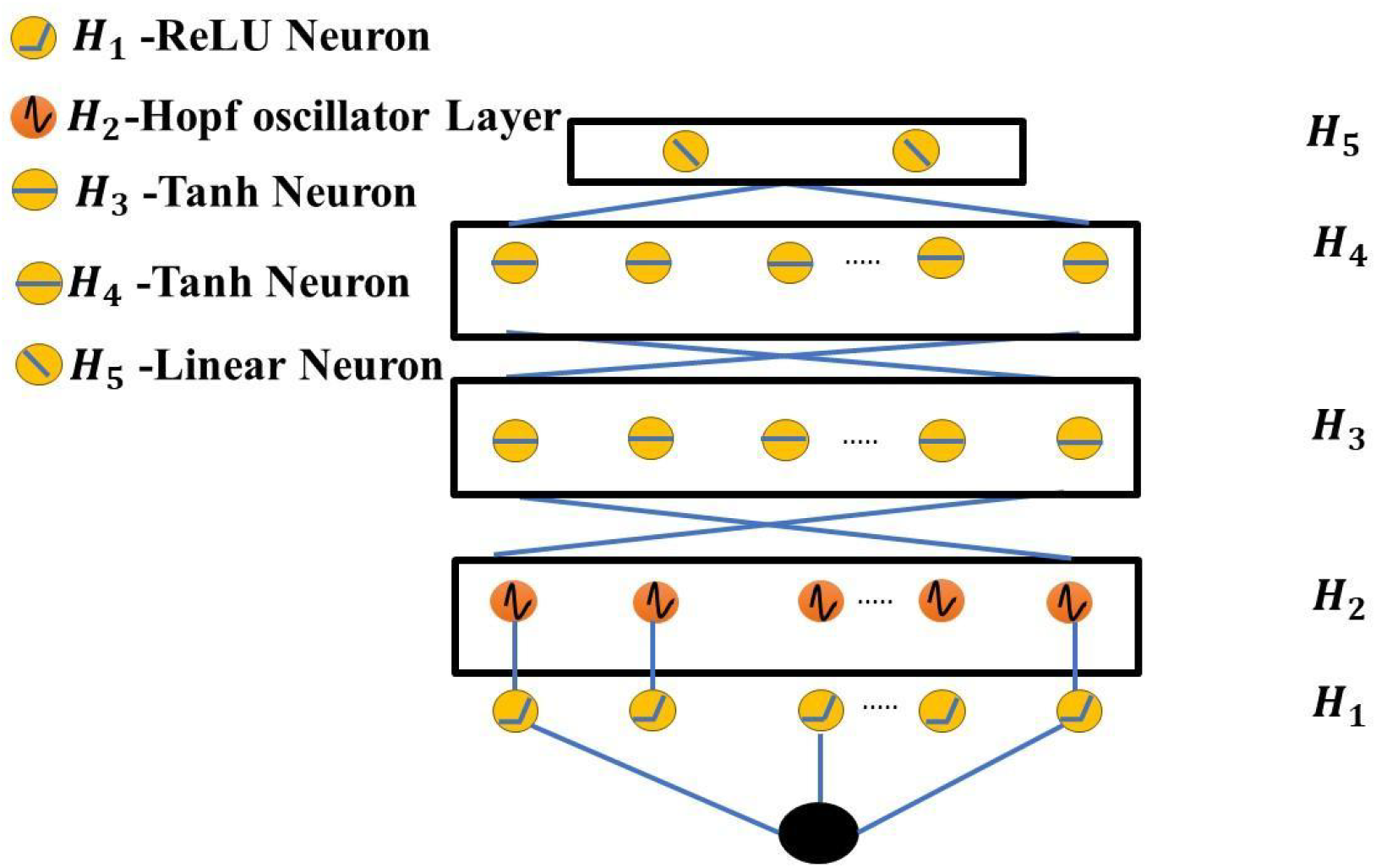
Model architecture of Deep Oscillatory Neural Network (DONN)

**Fig 2:**
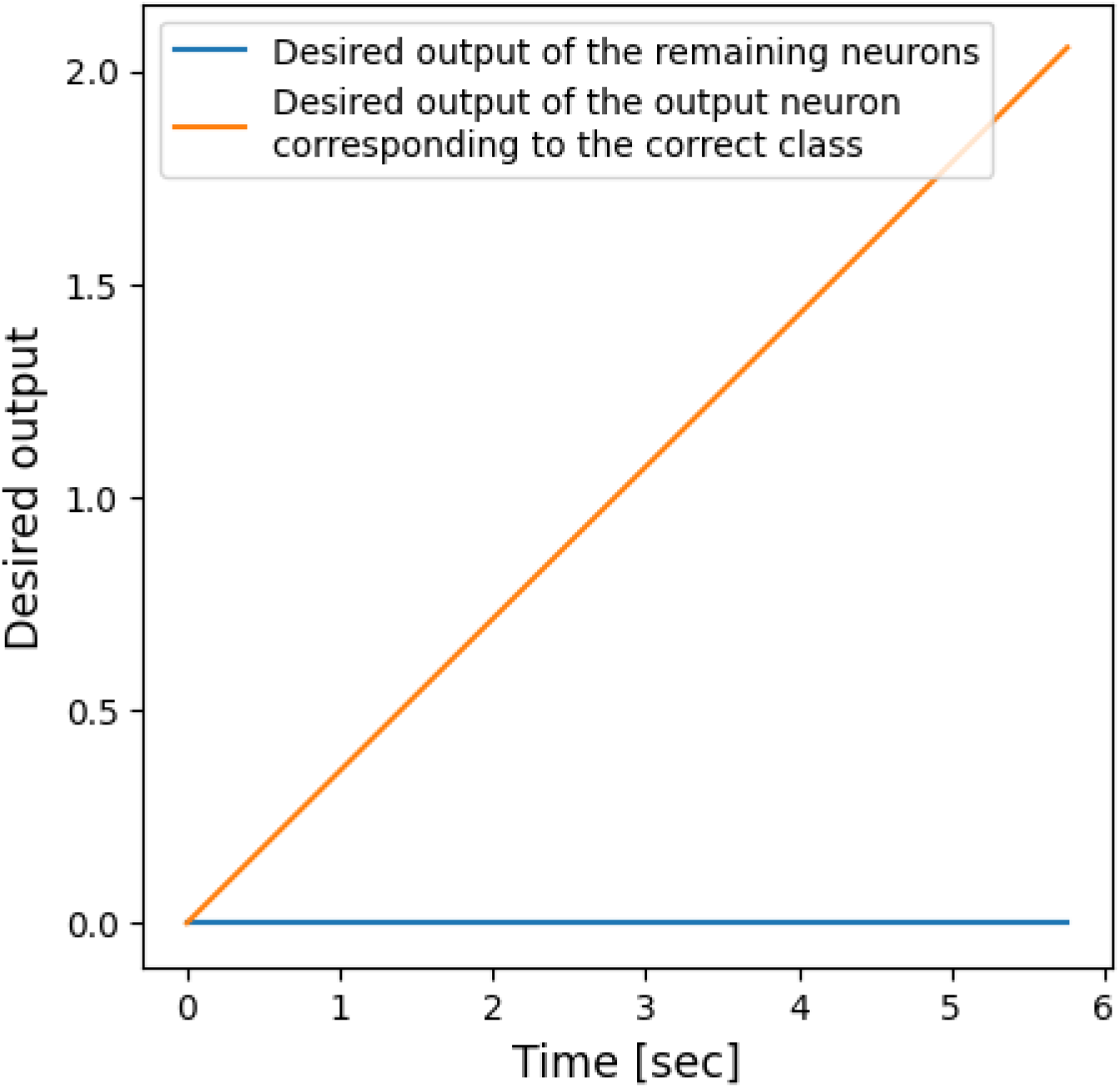
Desired ramp output for the output neuron connected to the correct neuron

The output of the H1 layer (ReLU layer) is presented as input to the H2 (layer of oscillators) as described in eqn (2.3). Note that the connection between H1 ReLU layer and H2 oscillatory layer is one-to-one whereas all other remaining connection stages are all-to-all connections.

The frequencies (ω_*j*_) of the oscillators in the oscillator layer, H2, are randomly initialized in the range that matches the input EEG signal frequencies. There are no lateral connections among the oscillators of the oscillator layer.

There is a complex-valued weight stage (*w*^*\**^) connecting the signal (*x*_*in*_) to H1 layer. Then it is passed through complex ReLU (crelu). It applies Rectified Linear Unit to both the real and imaginary part 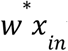 where *w*^*\**^ can be written as: 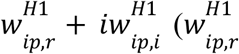 is the real part of connecting weight between the input to H1 layer, weight between input to H1 layer). 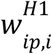 is the imaginary part of connecting

Relu is applied to the 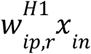 and 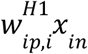

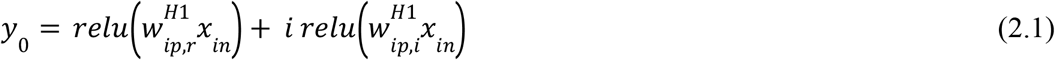

where normal relu() can be defined as:

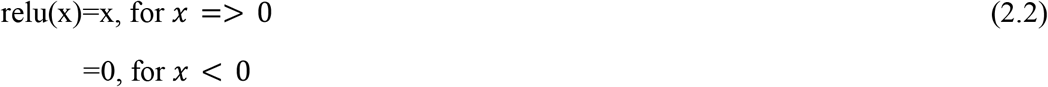

Equation (2.1) can be written as:

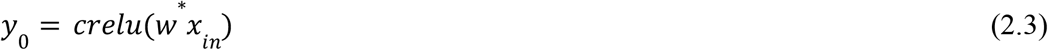

Furthermore, the H1 layer activations (*y*) are passed to the H2 layer (oscillatory layer) as an external input scaled by ξ. The oscillatory layer equations are described in eqns. (1.5-1.9). *z*_*k*_ is the complex valued output of H2 layer.

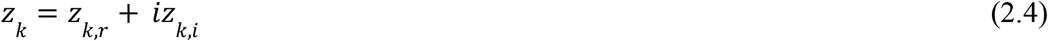

where *z*_*k,r*_, *z*_*k,i*_ are the real and imaginary part of H2 layer. Similarly,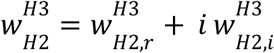

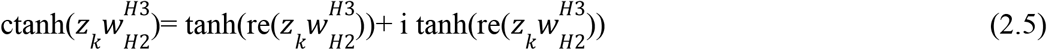

In the next two layers of the network, the neurons have tanh nonlinearity. Any number of hidden layers can be added successively to the network similar to deep networks depending on the problem at hand.

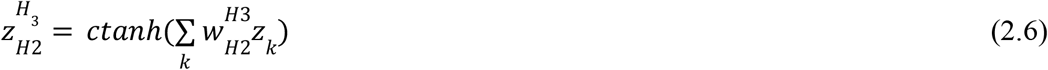

Next, the oscillatory outputs of the H2 layer (oscillatory layer) (eqns. 1.8-1.9.) are sent to the complex tanh layer (*H*_3_). Here also the connection weights 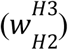 are complex-valued.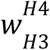 is the complex feedforward weight between *H*_3_ to *H*_4_ .

Therefore, forward propagation from *H*_3_ to *H*_4_ :

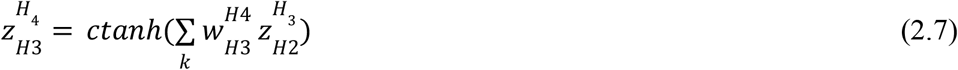

where 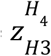 is the activation of *H*_4_ layer, and 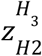 is the activation of *H*_3_ .

Any number of successive hidden layers can be added similarly.

The output layer has one linear neuron for the regression problem and as many linear neurons as there are classes for the classification problem. All the weights of the network are complex-valued.

The model shown in Fig. 2 is a generic one and can be used for both classification and prediction.

Hopf oscillator differential equations are solved using the forward Euler method with a fixed time step over the data length. All the simulations have been performed in Python3 using TensorFlow2 library. All the operations mentioned above are continuous and differentiable, and the network weights are updated using TensorFlow’s autograd.

### 3.3 DONN Model for EEG Time series classification

In this section, we describe the DONN model as a classifier model, which is the slightly modified version of the previously shown general DONN architecture (fig. 2). When the DONN model is used for classification, the number of output neurons, which are linear neurons, equals the number of classes, and all the hidden layer neurons are complex; at H5 layer we have real-valued neurons, which take only the real part of their inputs.

Signal classification is implemented by appropriately defining the desired output in a supervised learning scheme. In the case of vector classification, since it can be done in a single step, the desired class can be given in a “one-hot” representation. However, in the case of signal classification, since classification can only be done after examining the input signal for a finite duration, the desired output cannot be given in “one-hot” form throughout the duration of the presentation. Therefore, we set the desired output of the output neuron that corresponds to the correct class as a ramp signal, while the outputs of the remaining output neurons are zero throughout (fig. 3).

**Figure 3:**
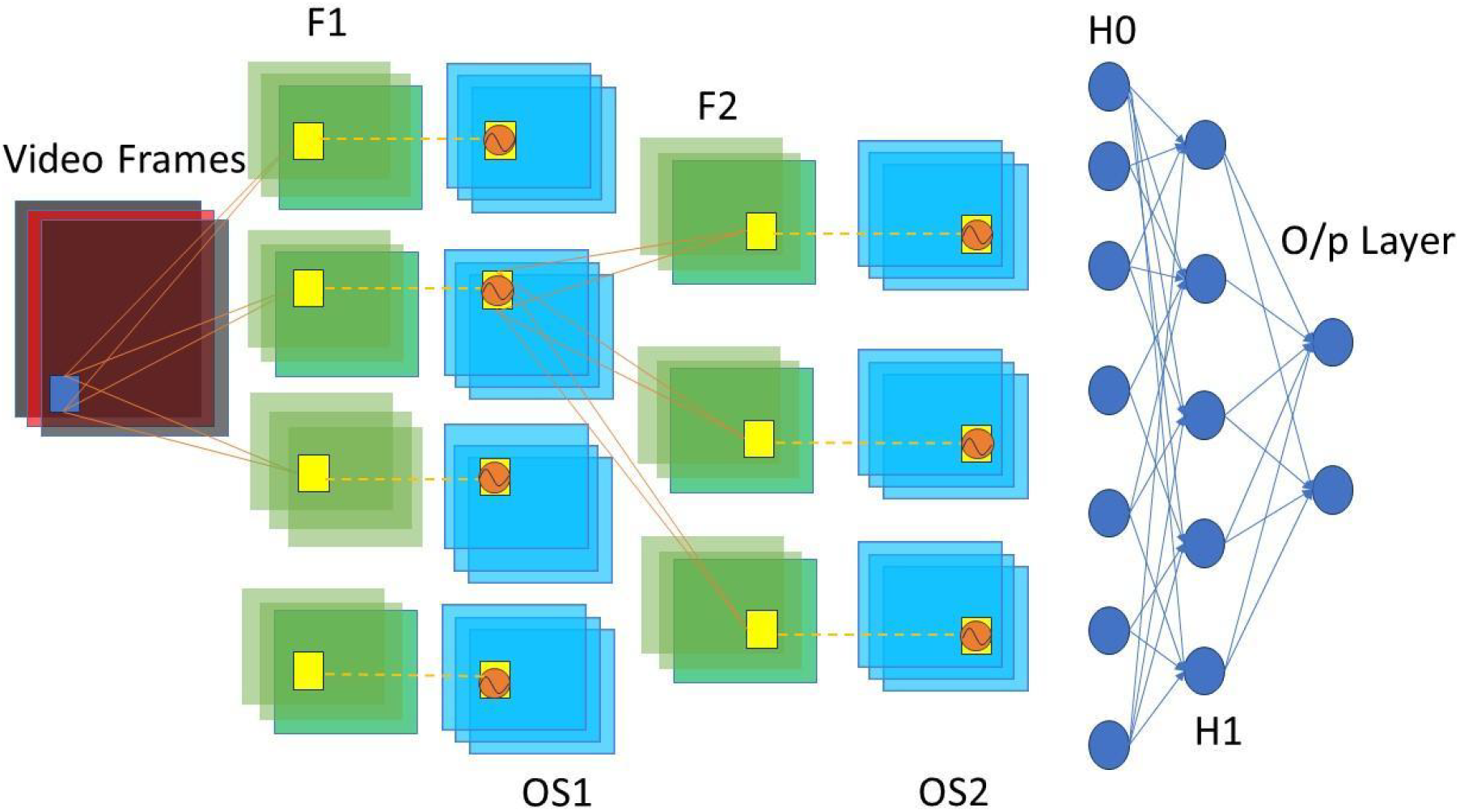
Model architecture of Oscillatory Convolution Neural Network, where F1-feature map layer-1, OS1-Oscialltory layer 1, F2-feature map layer 2, OS2-oscillatory layer 2, H0 – dense layer 0, H1-Dense layer 1, O/P – output layer.

### 3.4 Spatio-temporal Image Classification Using Oscillatory Convolutional Neural Network (OCNN)

In the DONN model, we presented a vector of EEG channels as input. However, we noted that the classification performance was not satisfactory as the number of input channels grew. Therefore, for multi-channel EEG we propose the *topoplot* as the input. Topoplot [20] is an excellent way to represent spatiotemporal activity of the human scalp from EEG which is elaborately discussed in section 3.4. To classify a time-varying image, we use an OCNN model for multi-channel EEG classification. The method for representing multichannel EEG data as the topoplot is described in Section 3.5.

In the original CNN, after the input layer, there exists an alternating series of convolutional and maxpooling layers, followed by a series of dense or fully connected layers. In OCNN, after the input image, there exists a series of alternating convolutional and oscillator layers, followed by a series of dense layers (fig. 4). The convolutional layers are comprised of sigmoidal neurons whereas the oscillator layers are made up of oscillator neurons. The dense layers have complex-valued sigmoidal neurons. These layers are described below.

**Figure 4:**
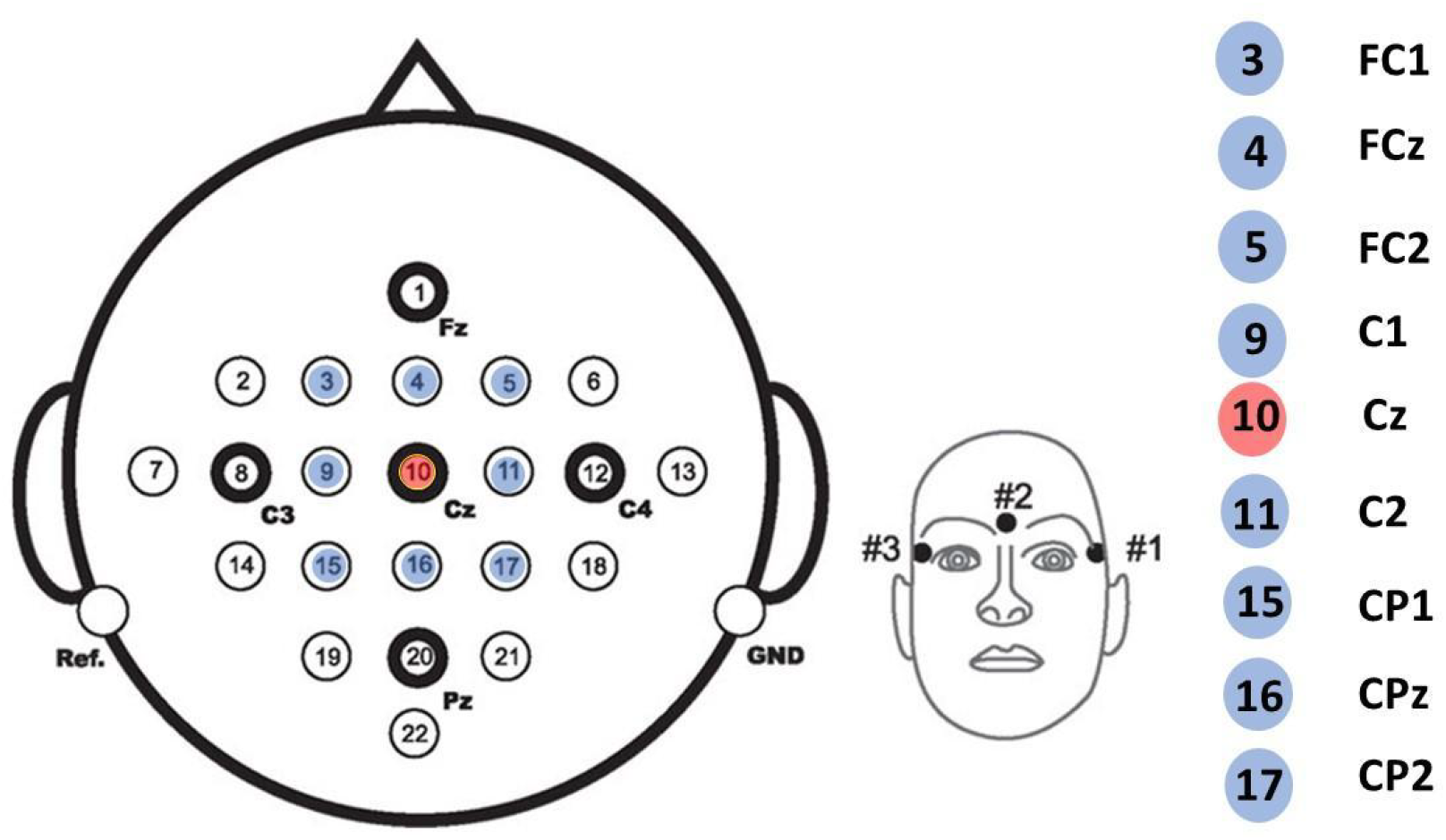
A graphical representation of target channel prediction from its neighboring channels

#### Input and Convolution layer (F1)

At first, we create topographic images in grayscale as discussed in section (3.5). Topographic images are then convolved (eqn. 3.1-3.2) as in the conventional CNN. The kernel weights are learnable; ReLU activation function is used. The resultant matrices from the convolution operation of the kernel when it moves across the height and width of the video frame are known as feature maps. The input video frame is of size (*I*_*d*_ X *I*_*d*_), where *I*_*d*_ is both the height and the width of each frame. So, the output volume of the feature map for kernel dimension (*N*_*f*_ *X N*_*f*_) is calculated using the following formula:

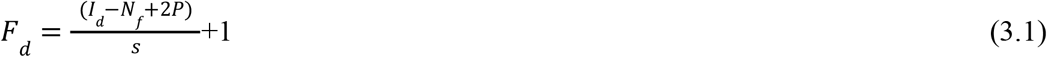

where *I*_*d*_ is the size of the input image, P is the padding, and s is the stride length.

Then each pixel value of the feature map is fed as the Hopf oscillator’s external input (*I*(*t*)) at each time instant. Therefore, the oscillatory layer’s dimension is equal to the feature map dimension. This combination of convolutional oscillatory blocks can be stacked successively similar to convolutional neural networks.

The size of each feature map is given as:

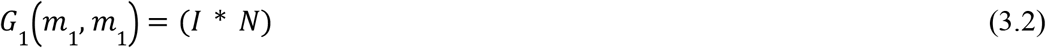

where, *G*_1_(*m*_1_, *m*_1_)is the feature map of size m1, *I* is the input image, N is kernel of size (*N*_*f*_ *X N*_*f*_ *X F*_*N*1_) (*F*_*N*1_ is number of kernels).

#### Oscillatory Layer (OS1)

The size of OS1 (oscillators layer) is the same as F1 layer. Therefore, the number of oscillators present in the OS1 layer is (*G*_1_ (*m*_1_, *m*_1_), *f*_*N*1_). Here, the connections between the feature map and the oscillator layer are one-to-one. Hopf oscillators’ frequencies are randomly initialized based on the FPS (frame per second) of the input video frames. For the oscillators, the state variable dynamics equation with the forced external signal is described by eqns. (1.5-1.9). The output from the oscillatory layer at any time instant is of dimension (*G*_1_ (*m*_1_, *m*_1_), *f*_*N*1_).

From the OS1 (1^st^ oscillatory layer) we again apply a kernel of size (*N*_*f*2_, *N*_*f*2_) and the number of filters are *F* . The size of F2 (feature map layer 2) is also calculated by using eqn. 3.2. (*f*_*N*2’_, *G*_2_ (*m*_2_, *m*_2_)). After that, OS2 (oscillatory layer) has the same size as F2. Here also the feature map’s output values are also fed into the oscillator’s external signal as described by eqns. (1.5-1.9). After OS2 (2^nd^ oscillatory layer), we flatten the network (H0 layer). Thus, the size of H0 layer is *f*_*N*2_ * *G*_2_ (*m*_2_, *m*_2_).

#### Dense layer(s)

Then the H0 layer’s output is multiplied with complex weights 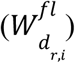 and sent to H1 layer, where,

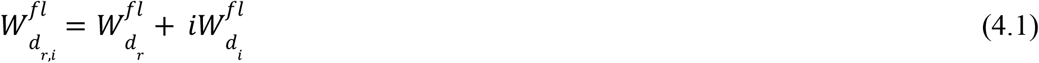

The output of H1 (1^st^ hidden dense layer) is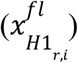 :

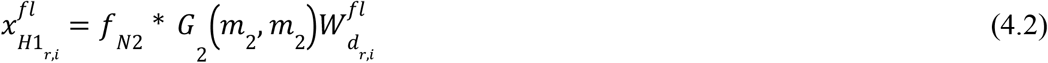

After that H1’s output is fed into the output layer having 2 linear neurons for the two class classification problem.

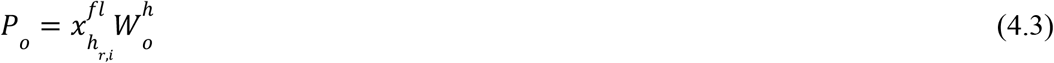

### 3.5 EEG Topographical Mapping: constructing the Topoplot

The topoplot is a convenient way to visualize EEG data over the electrode space [20]. It depicts the spatial distribution of voltage activity over the scalp surface. It is a 2-D circular view, created based on interpolation on a fine cartesian grid. It is also a useful measure for data cleaning and data inspection and a convenient way to visualize brain network dynamics. It is also used for 2-D visualization of power values of EEG data. Data vector is a single-column vector that has the corresponding data. The topoplot function available in MNE-python is designed for visualizing power values in the default scale -1 to +1 mapped to the intensity of the color at a particular region. It plots a topographic map of a scalp data field in a 2-D circular view using interpolation on a fine Cartesian grid. In the topomap function, we have to select a specific montage. Currently we chose the standard ‘biosemi-64’ montage which takes the standard electrode’s cartesian coordinates.

## 4. Results

### 4.1 Nearest channel Data prediction on motor imagery Dataset

The data for the EEG channel prediction study is obtained from BCI 2A experiment discussed in section 2.1.5. The 8 channels/electrodes used on the input side are: fc1, fcz, fc2, c1, c2, cp1, cpz, cp2, whereas the Cz electrode is used for prediction (Fig. 5). Note that fig. 5 shows the graphical orientation of 10-20 electrodes [21]. The duration of each sample is 4 s (1000 time points). During training we use 9 channels of 1000 data points for each sample. We took ‘Cz’ channel for prediction which is the centrally located over other 8 channels (3,4,5,9,11,15,16,17) marked in fig. 5. The validation set is 20% of training data. All the model parameters are given in Table 1 below. The model prediction (testing) performance is calculated using RMSE error. Average RMSE is calculated over all the testing samples. The average RMSE as well as training and validation loss is shown in Table 2. All the training and testing is done on individual subject basis. Model predicted EEG Data (Cz channel) for subject 2 is shown in fig. 6.

**Table 1:** Model parameters for nearest channel prediction using DONN.

|  |  |
| --- | --- |
| Model parameters |  |
| Input signal size(training) | 400,1000 |
| Testing sample | 100,1000 |
| H1 Layer | ReLU neuron, units-50 |
| H2 | Hopf oscillators , units-50 |
| H3 | Tanh neuron, units-40 |
| H4 | Tanh neuron, units-30 |
| H5 | Linear neuron, units 1 |
| Hopf oscillator frequencies | [0.1-10 Hz] |
| Oscillator external input scaling factor ($\epsilon$) | 4 |
| Oscillator's $\mu, \beta$ | 1,-4 |
| Learning rate | 0.001 |
| optimizer | adam |
| Number of epochs and batch | 100, 4 |

**Table 2:**
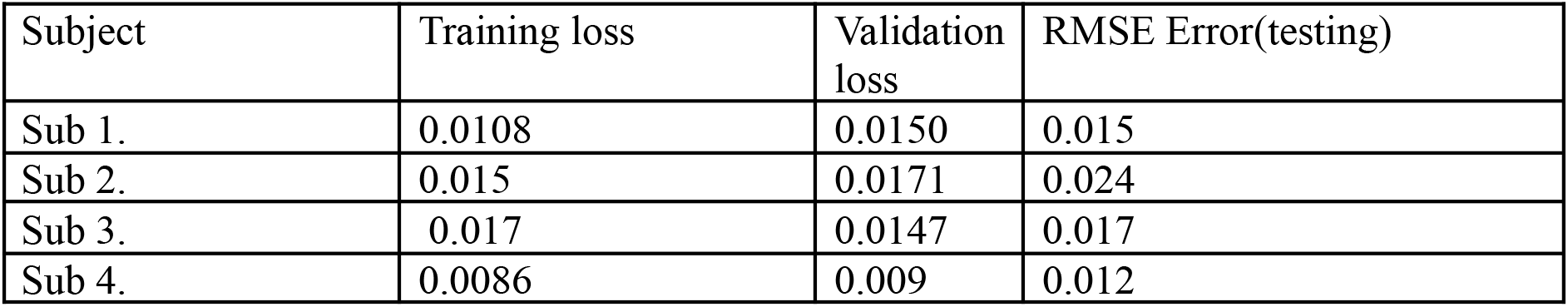
Testing -training error all 4 subjects from BCI Dataset.

**Fig 5:**
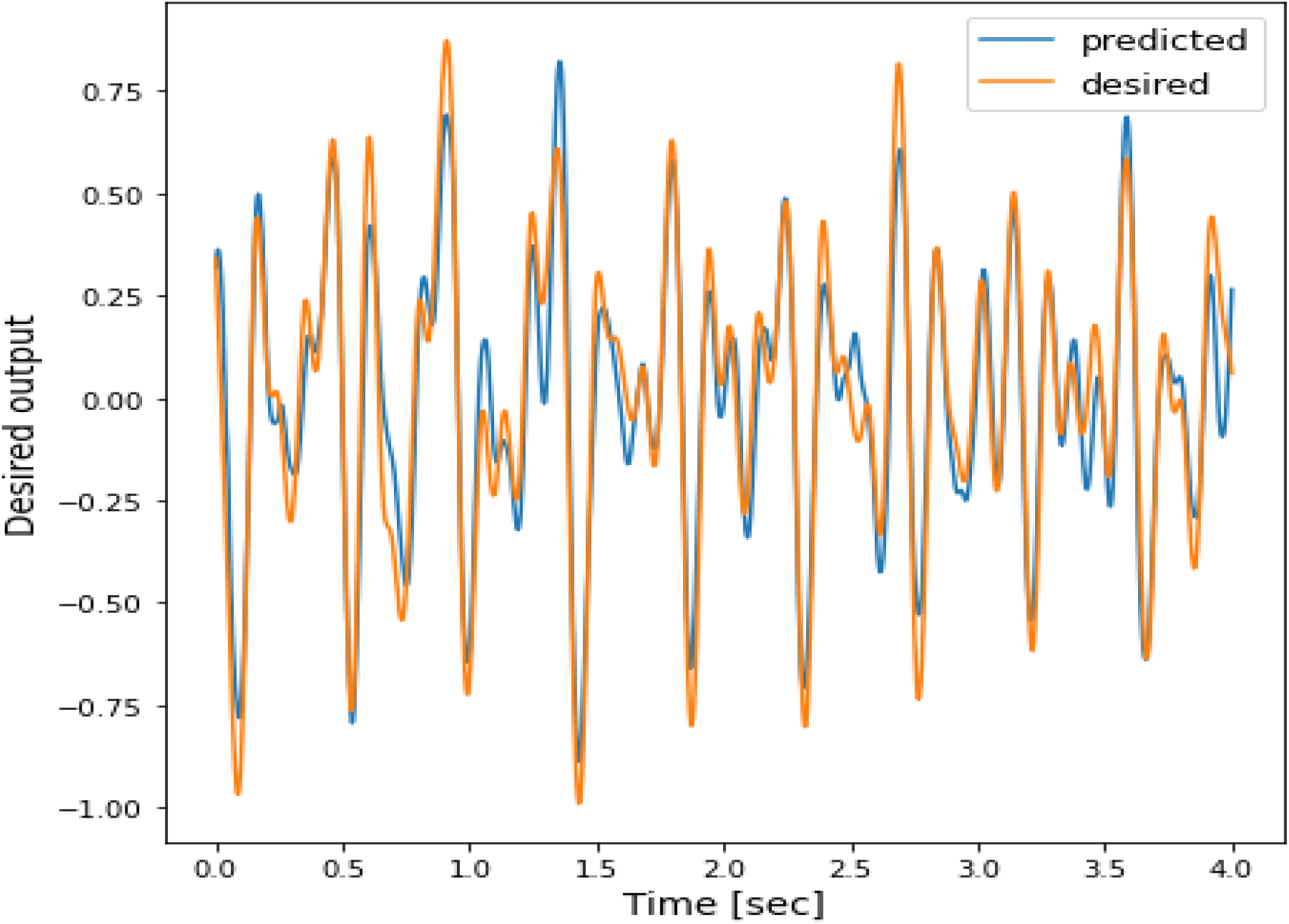
EEG Prediction for the central channel (cz) based on the following neighboring channels: fc1, fcz, fc2, c1, c2, cp1, cpz, cp2.

**Fig 6:**
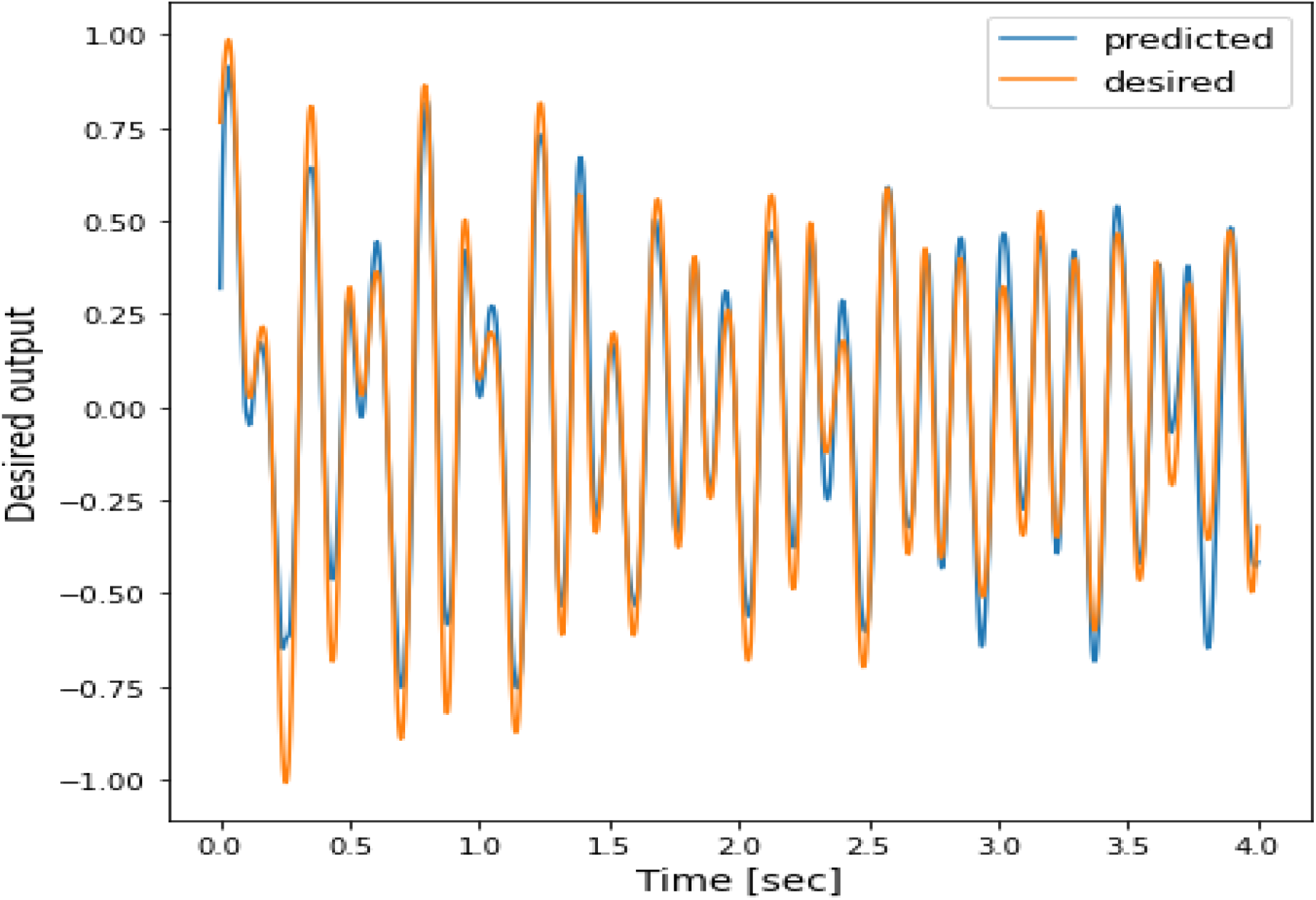
EEG Prediction for the central channel (cz) based on the following neighboring channels : fc1, fcz, fc2, c1, c2, cp1, cpz, cp2 for subject 5.

**Fig 7:**
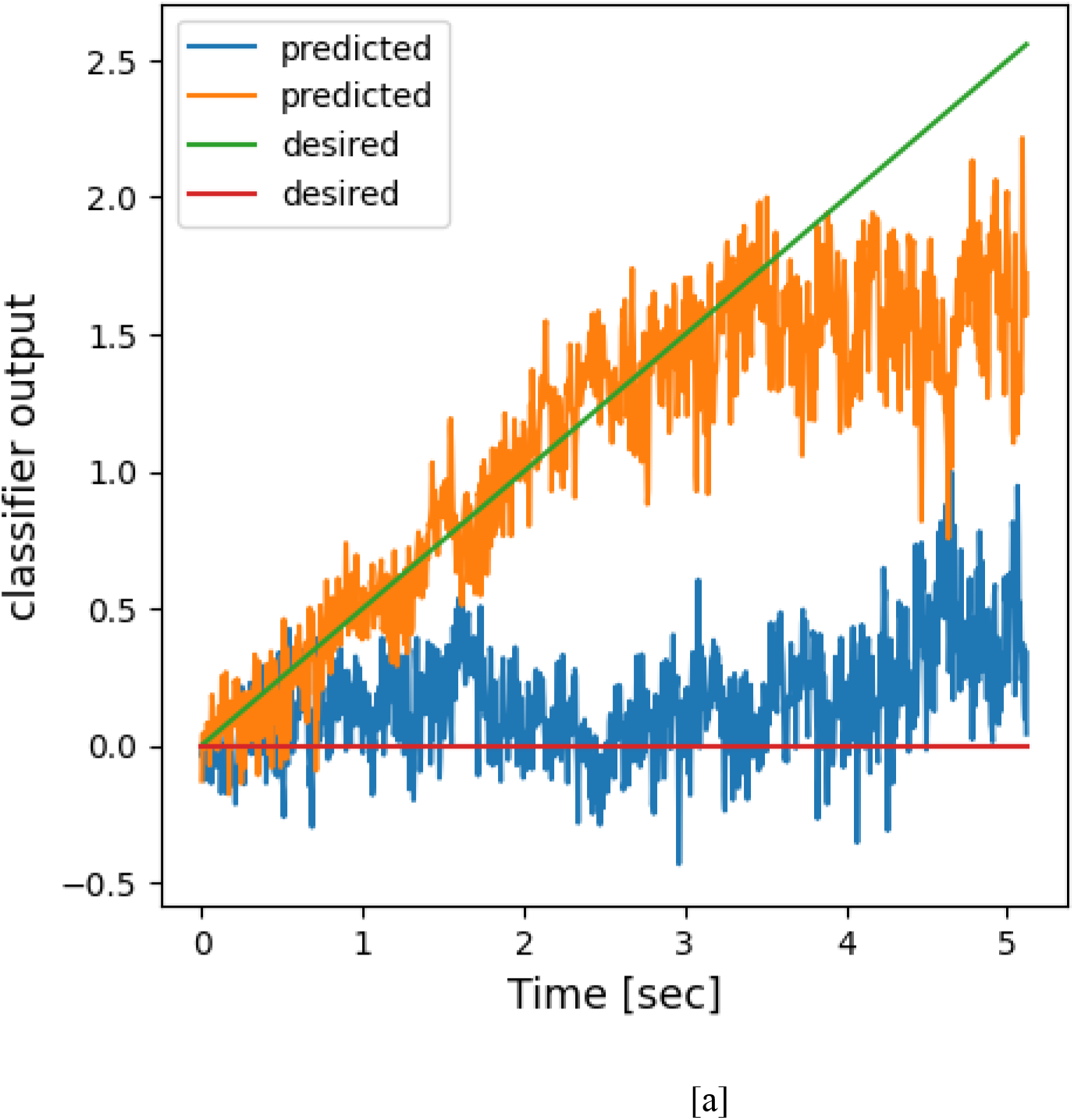
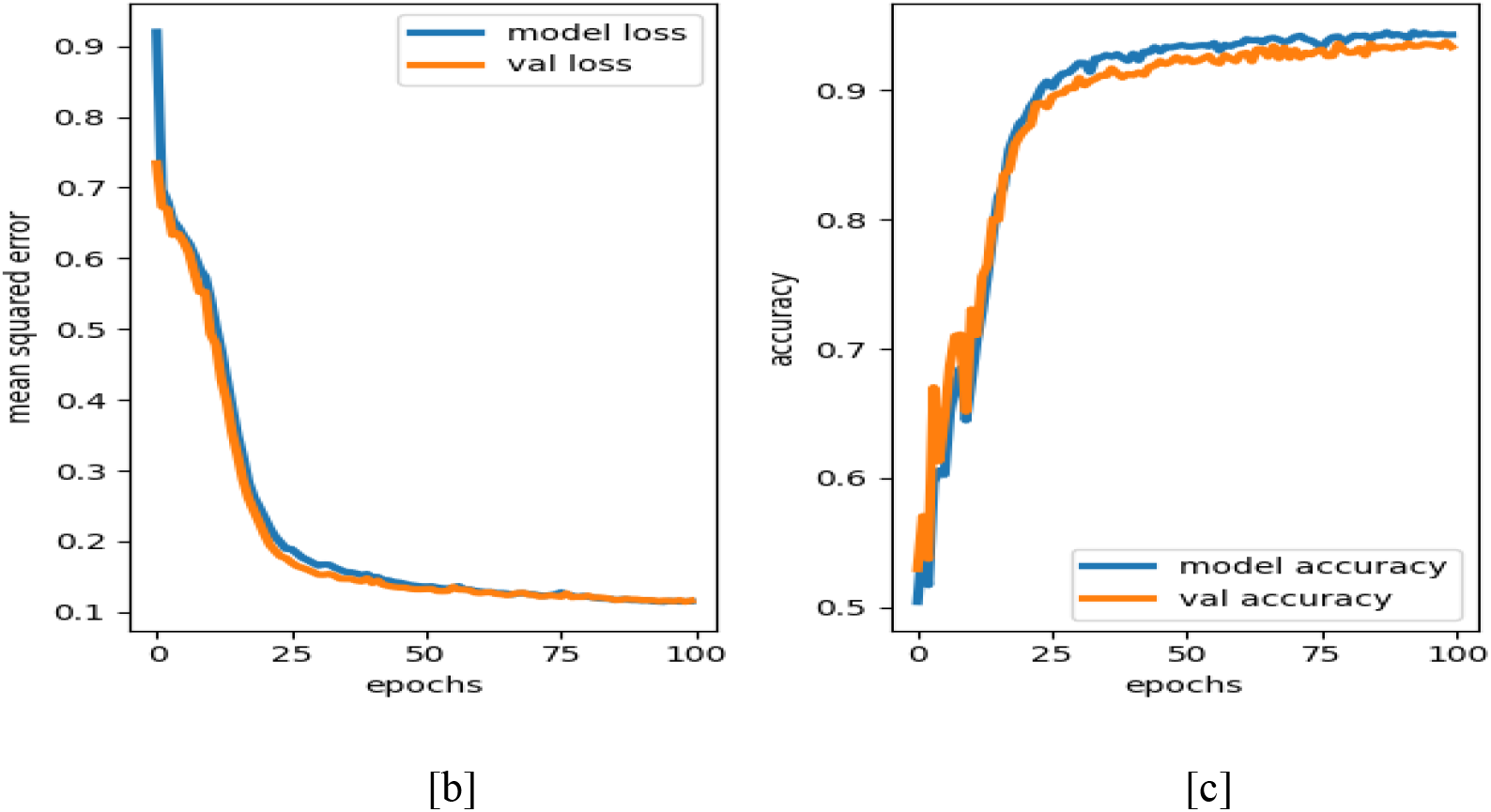
[a]: The target output and the desired output (during testing) plot for pre-ictal Vs. ictal classification, [b-c]: RMSE-Validation accuracy for pre-ictal vs. ictal EEG classification

Another application of DONN we explored here, where training and testing is done on multiple subjects basis. We used four subjects’ data for training but testing is done on 5^th^ subject. Same model parameters are used as mentioned in Table 1. The model obtained a testing RMSE of 0.013. Model Predicted signal of Cz channel from 5^th^ subject along with the desired testing signal is shown in fig. 6.

### 4.2 Classification based on single channel EEG

For the three classification studies described below, we use single channel EEG data from IITD epilepsy dataset, BONN Dataset and Physionet Sleep dataset (supplementary file).

#### 4.2.1 IIT Delhi (IITD) Epilepsy dataset

Using the data sets described in section (2.1.1), we applied DONN models for the following two and three class problems: (a): pre-ictal vs ictal (b): inter-ictal vs ictal (c) pre-ictal vs inter-ictal vs ictal.

In the case of pre-ictal vs. ictal data, the length of EEG data is 1024 samples. 80% of the data is used for training and the rest is used for testing. From H1 to H2 layer one-to-one connections are used and other feedforward connections are all-to-all. A detailed description of model parameters is shown in table 3.

**Table 3:** The model parameters for IIT D epilepsy data classification using DONN.

|  |  |
| --- | --- |
| Model parameters |  |
| Input signal size | 100,1024(2-class problem)/150,1024(3-class problem) |
| H1 Layer | ReLU neuron, units-50 |
| H2 | Hopf oscillators , units-50 |
| H3 | Tanh neuron, units-40 |
| H4 | Tanh neuron, units-30 |
| H5 | Linear neuron, units-2(2 class problem)/3(for 3 class problem) |
| Hopf oscillator frequencies | [0.1-70 Hz] |
| Learning rate | 0.01 |
| optimizer | adam |
| Oscillator external input scaling factor ($\epsilon$) | 4 |

In the case of the inter-ictal vs. ictal classification also, we used the same architecture (fig. 2). In this three-class problem, 80% of the entire data is used for training, and 20% is used for testing. The other network parameters are the same as the previously explained 2-class problem (table 3). Our model performance is compared against previously reported performances in Table 4. Model performance is shown in fig. (8(a-c)).

**Table 4:** Performance comparison of the proposed DONN model with previously published results.

| Methods | Result |
| --- | --- |
| [22] wavelet coefficient, and General Regression neural Network, Non-seizure Vs. Seizure | 97.93±0.217 |
| [23] Wavelet coefficient at different sub band of EEG signal(Energy, Entropy and Standard deviation feature) & probabilistic Neural Network., Normal vs. Epileptic | 99.33% (with energy feature, time per run is very less) |
| [73] EEG signal(0 to 60 Hz, DCT filter bank, EEG band extraction, hurst component & ARMA feature , SVM , Inter-Ictal Vs. Ictal Pre Ictal Vs. Ictal classifier) | 97%<br>74.60% |
| Deep oscillatory neural network (proposed), 2class, Pre-ictal vs ictal | **100%** |
| Deep oscillatory neural network (proposed), 2 class, Inter-Ictal vs. Ictal | **95%** |
| Deep oscillatory neural network (proposed), 3 class, Pre-ictal vs. inter ictal vs ictal | **96.67%** |

**Figure 8.**
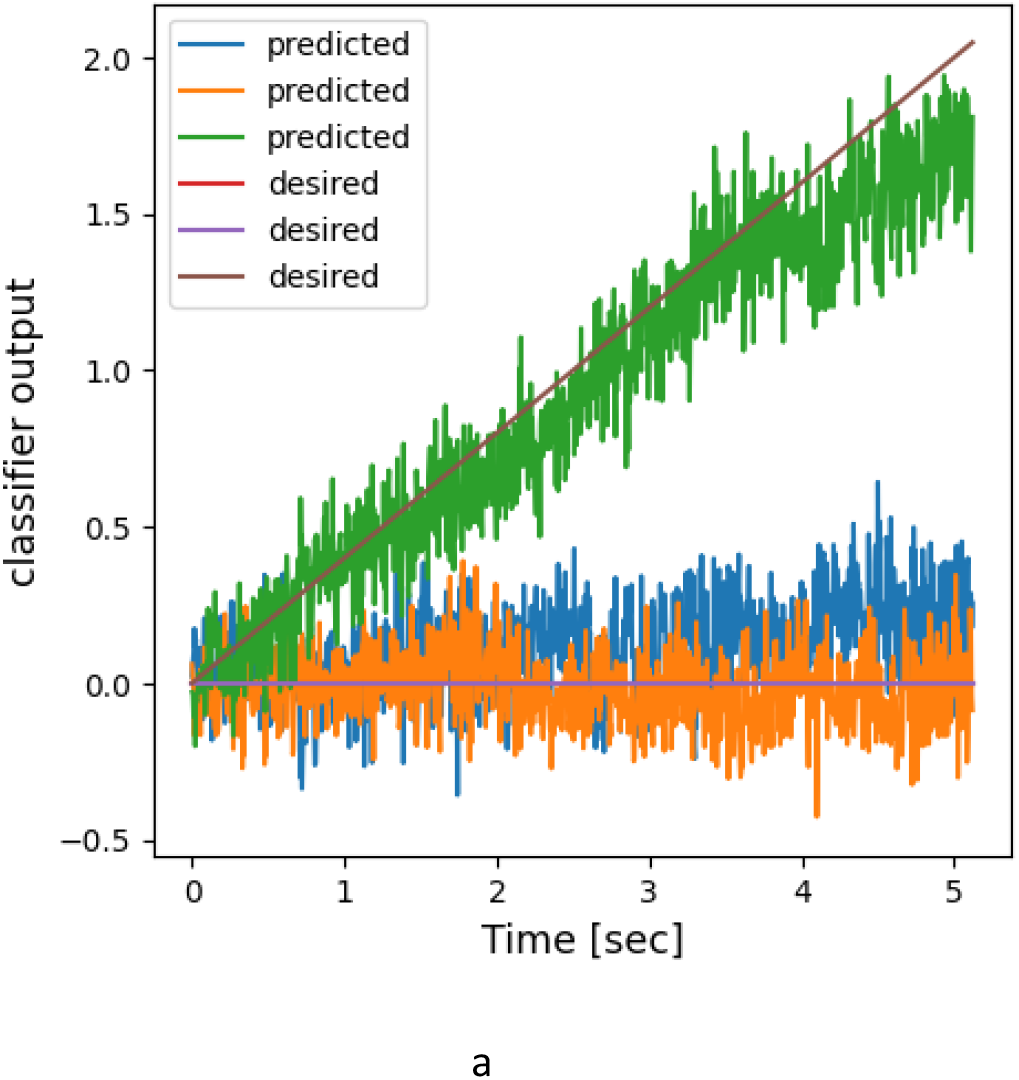
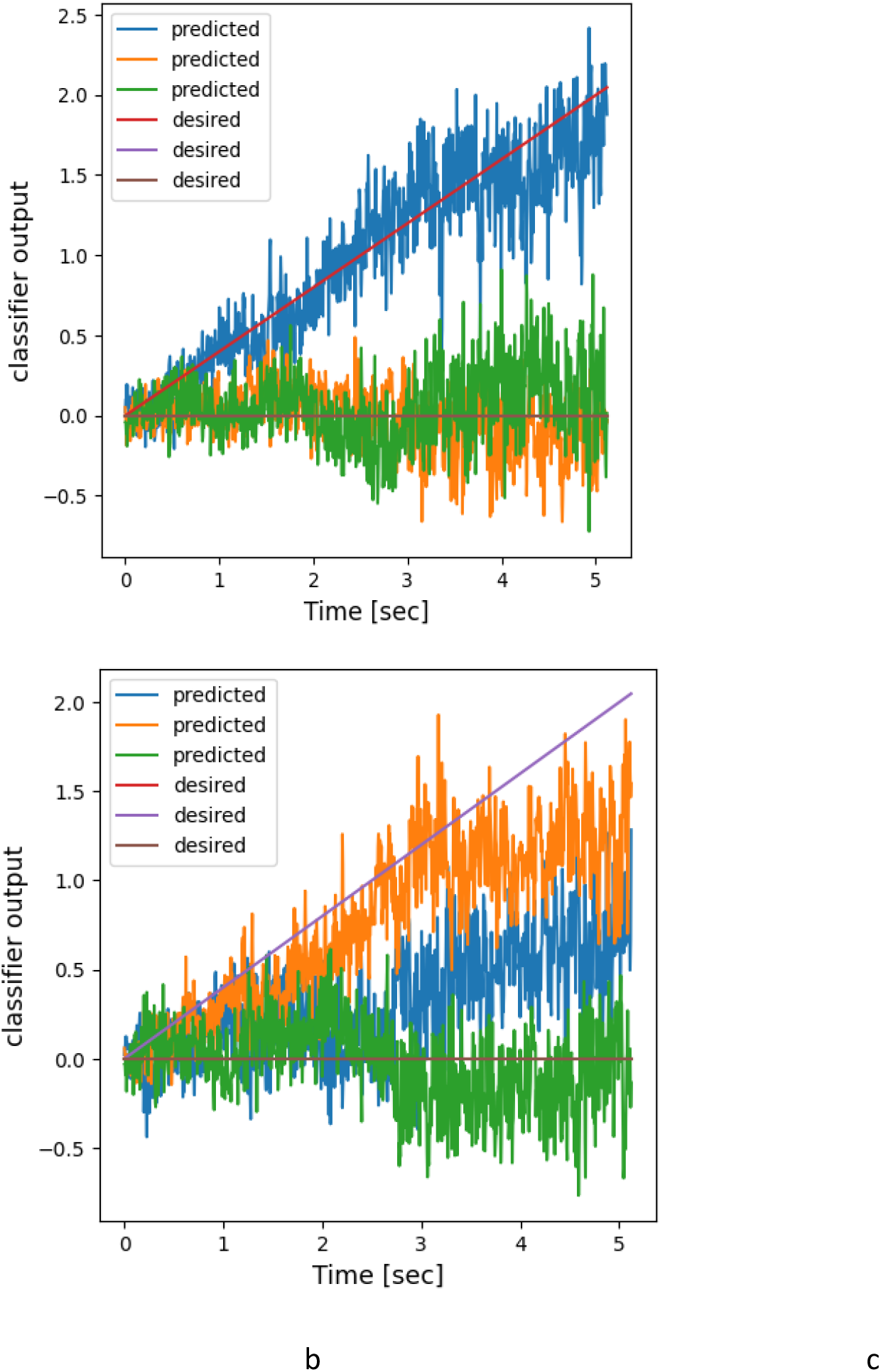
(a-c): testing output (target and model predicted) for the 3-class problem (pre-ictal vs. inter ictal vs ictal)

**Figure 9.**
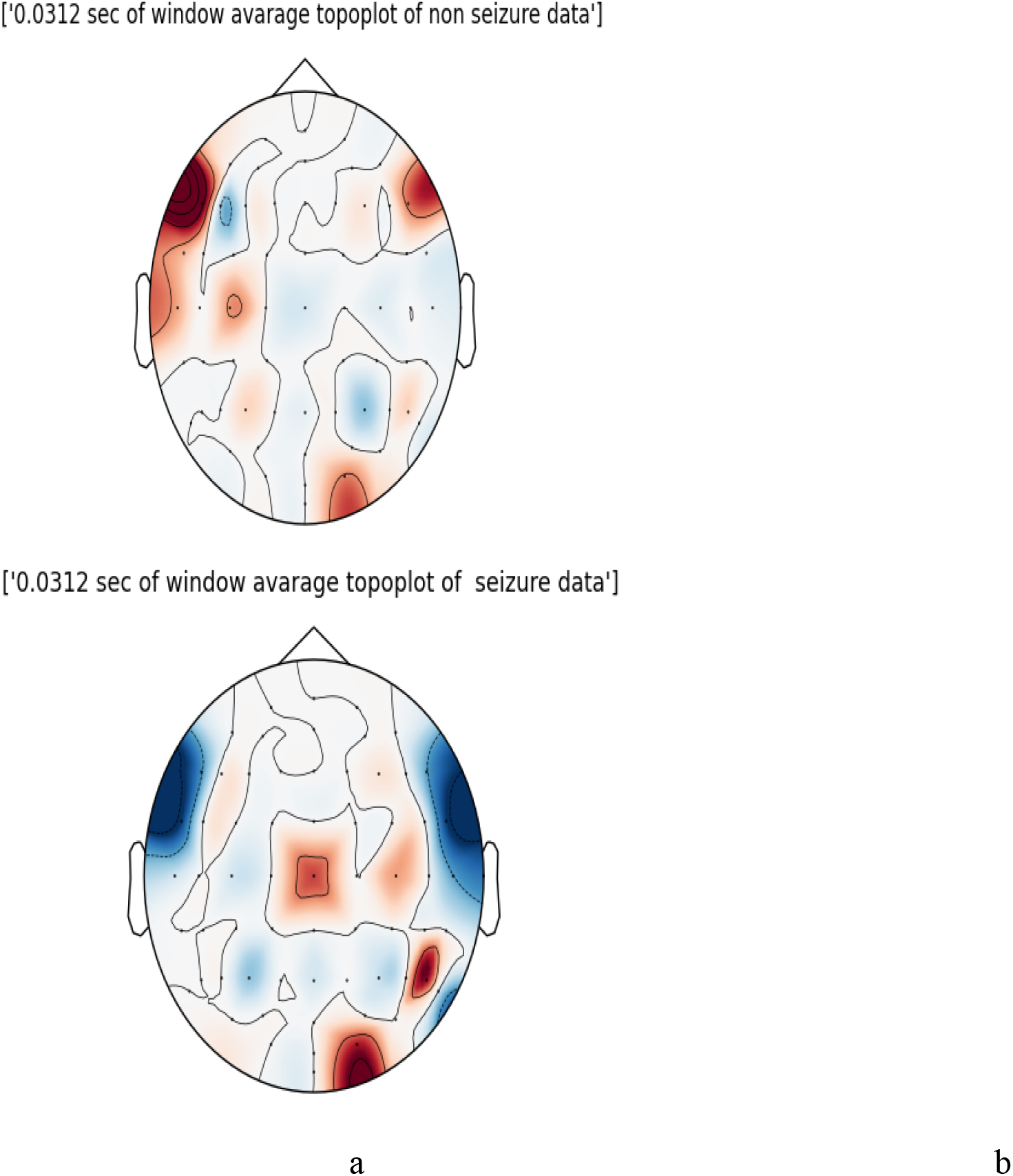
Topomaps of non-seizure (a) and seizure (b) data.

#### 4.2.2 BONN Dataset

As described in Section (2.1.2), BONN dataset consists of all three classes (healthy eye open-A, healthy eye closed-B, inter ictal-C, D, and ictal-E). Here we have done five different classifications including five 2-class classification problems and one 3-class classification problem. Here we used 1000 sample data points for each sample. For all the cases of 2-class problems, the model parameters are described in section (table 5). In case of a 3-class problem (A-C-E), there is a slight change in the number of oscillators used to enhance testing accuracy. In that case, we used 80 oscillators. A comparison of some previously reported literature on BONN data classification with our proposed model performance is shown in Table 6.

**Table 5:** The model parameter of DONN applied to BONN data classification.

|  |  |
| --- | --- |
| Model parameters |  |
| Input signal size | 800,1000(2-class problem)/1200,1000(3-class problem) |
| H1 Layer | RelU neuron, units-50(2-class)/80(3-class) |
| H2 | Hopf oscillators, units-80 |
| H3 | Tanh neuron, units-40 |
| H4 | Tanh neuron, units-30 |
| H5 | Linear neuron, units-2(2 class problem)/3(for 3 class problem) |
| Hopf oscillator frequencies | [0.1-70 Hz] |
| Learning rate | 0.01 |
| optimizer | adam |

**Table 6:**
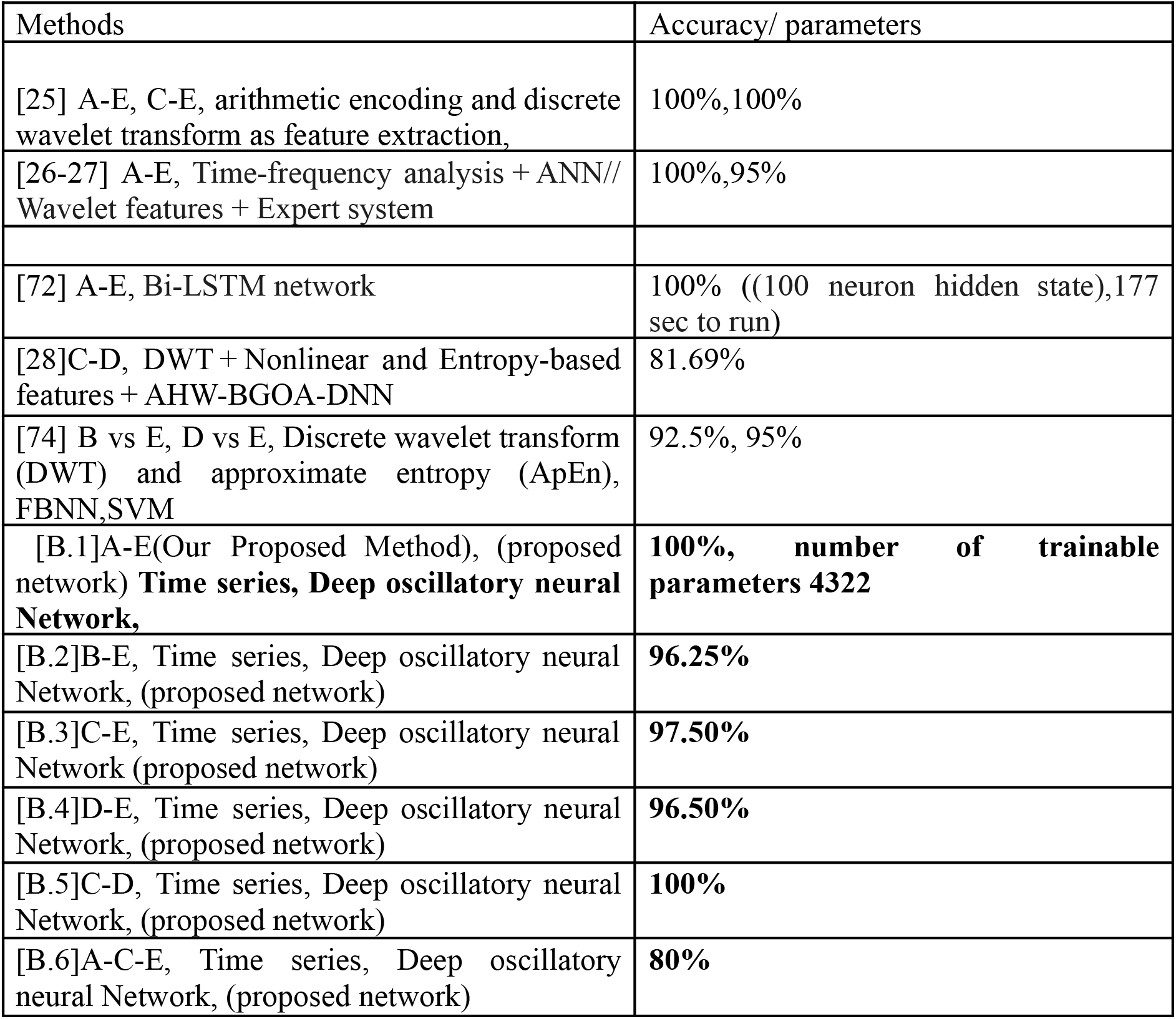
Performance comparison of the proposed DONN model with previously published results for the BONN Dataset.

**Table 7:** Describing the seizure events for CHB -01.

| Seizure data | Number of seizures | Seizure onset – seizure offset = Duration(sec) |
| --- | --- | --- |
| CHB01-03 | 1 | 2996-3036=40 |
| CHB01-04 | 1 | 1467-1494=27 |
| CHB01-15 | 1 | 1732-1772=40 |
| CHB01-16 | 1 | 1015-1066=51 |
| CHB01-18 | 1 | 1720-1810=90 |
| CHB01-21 | 1 | 327-420=93 |
| CHB01-26 | 1 | 1862-1963=101 |

#### 4.2.3 Classification of CHB-MIT Dataset in spatio-temporal form using Oscillatory Convolutional Neural Network (OCNN)

We performed 2 class classification with epileptic seizure and non-seizure data on CHB-MIT dataset, described in section 2.1.3. During data preparation, seizure data is extracted as mentioned in the description summary file and non-seizure data is extracted from the rest of the dataset. Firstly, we extract the entire epileptic segment from the subject’s data and create 2 s samples for every single subject, the number of time steps is 512 (sampling frequency is 256Hz) and the number of channels (electrodes) is 19. Then we applied non-overlapping window averaging over the duration of 0.015 s. Therefore, the final data dimension of single sample becomes 64x19, which is used to generate topographical maps. Some subjects contain multiple seizure instances and we made sure each sample from the individual is either seizure or normal instance.

After data segmentation and preprocessing we calculate topoplot using MNE toolbox in python [41]. This topoplot is calculated at single time points for all 19 electrodes (fig 10(a-b)). In MNE, using MNE viz. plot_topomap function, we have created spatio-temporal images of brain activity at different time instants. As the number of time points are 64, topoplot creates a video with 64 frames. After generating images for each frame, we normalized them in the range of [0 to 1] and resized the images to (50*50) size. Thus, the entire size of the video is 64x50x50 for every sample. These video sequences are fed to the OCNN networks. The architecture of the OCNN used in this study is defined in Table 8.

**Table 8:**
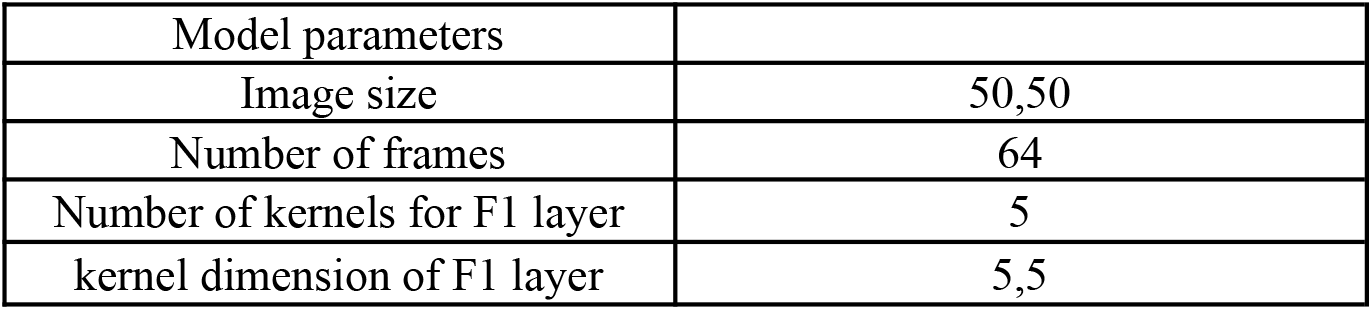
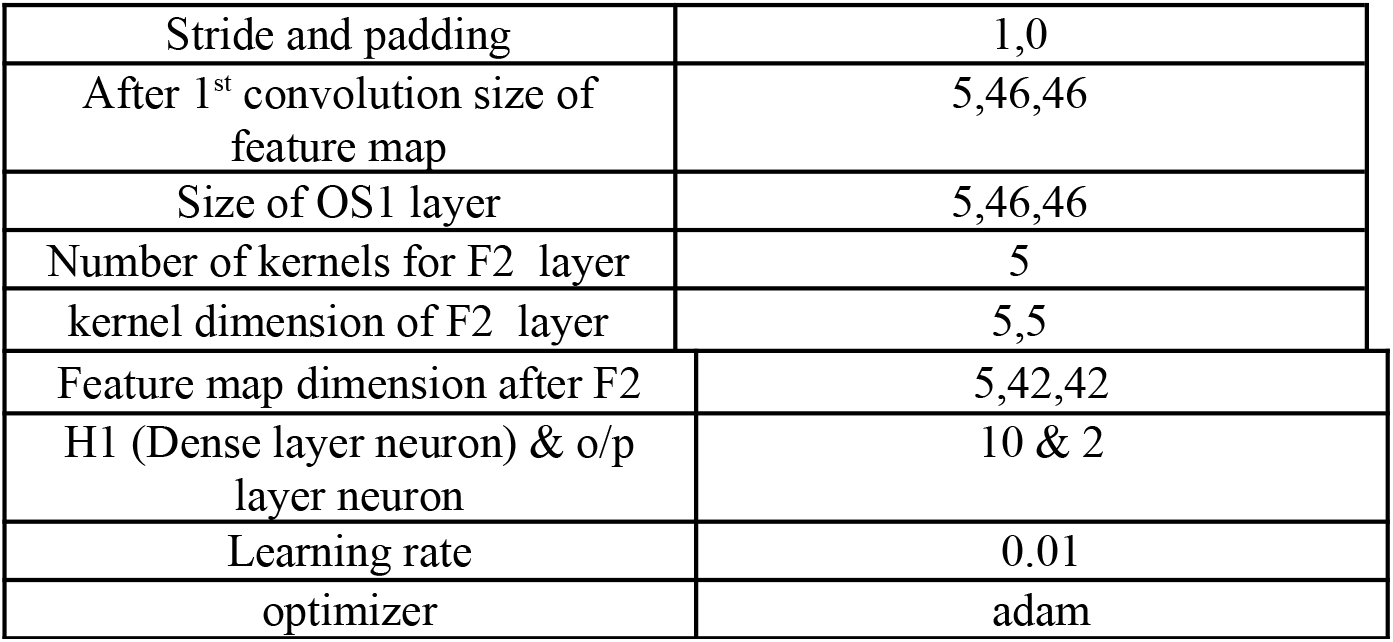
Details of the OCNN architecture used for CHB-MIT data classification.

| Model parameters |  |
| --- | --- |
| Image size | 50,50 |
| Number of frames | 64 |
| Number of kernels for F1 layer | 5 |
| kernel dimension of F1 layer | 5,5 |
| Stride and padding | 1,0 |
| After 1<sup>st</sup> convolution size of feature map | 5,46,46 |
| Size of OS1 layer | 5,46,46 |
| Number of kernels for F2 layer | 5 |
| kernel dimension of F2 layer | 5,5 |
| Feature map dimension after F2 | 5,42,42 |
| H1 (Dense layer neuron) & o/p layer neuron | 10 & 2 |
| Learning rate | 0.01 |
| optimizer | adam |

**Fig 10:**
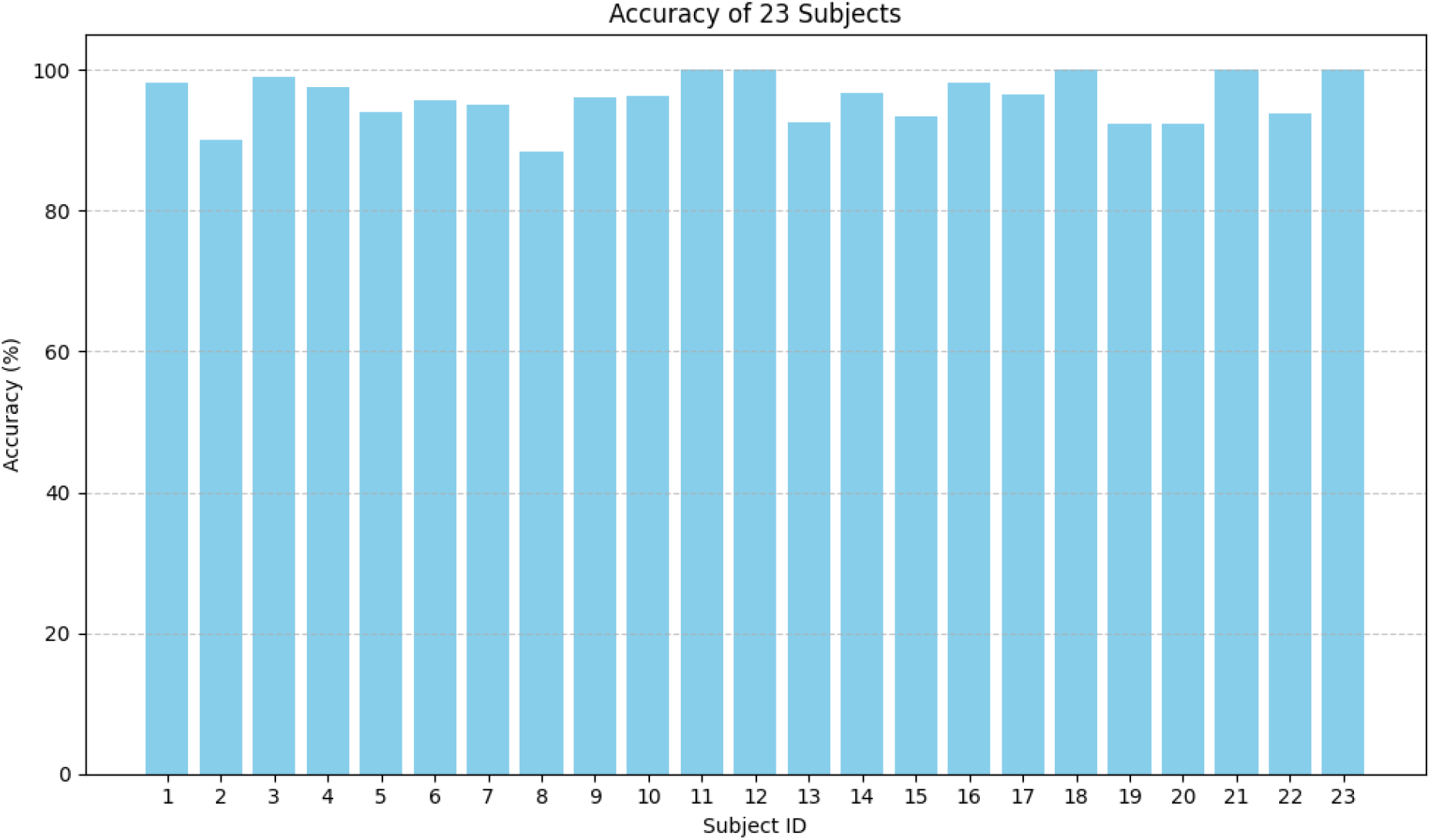
Our proposed model performance on CHB-MIT for all 23 subjects (accuracy reported per subject). Average accuracy over all 23 subject is 95.98%.

**Figure 11:**
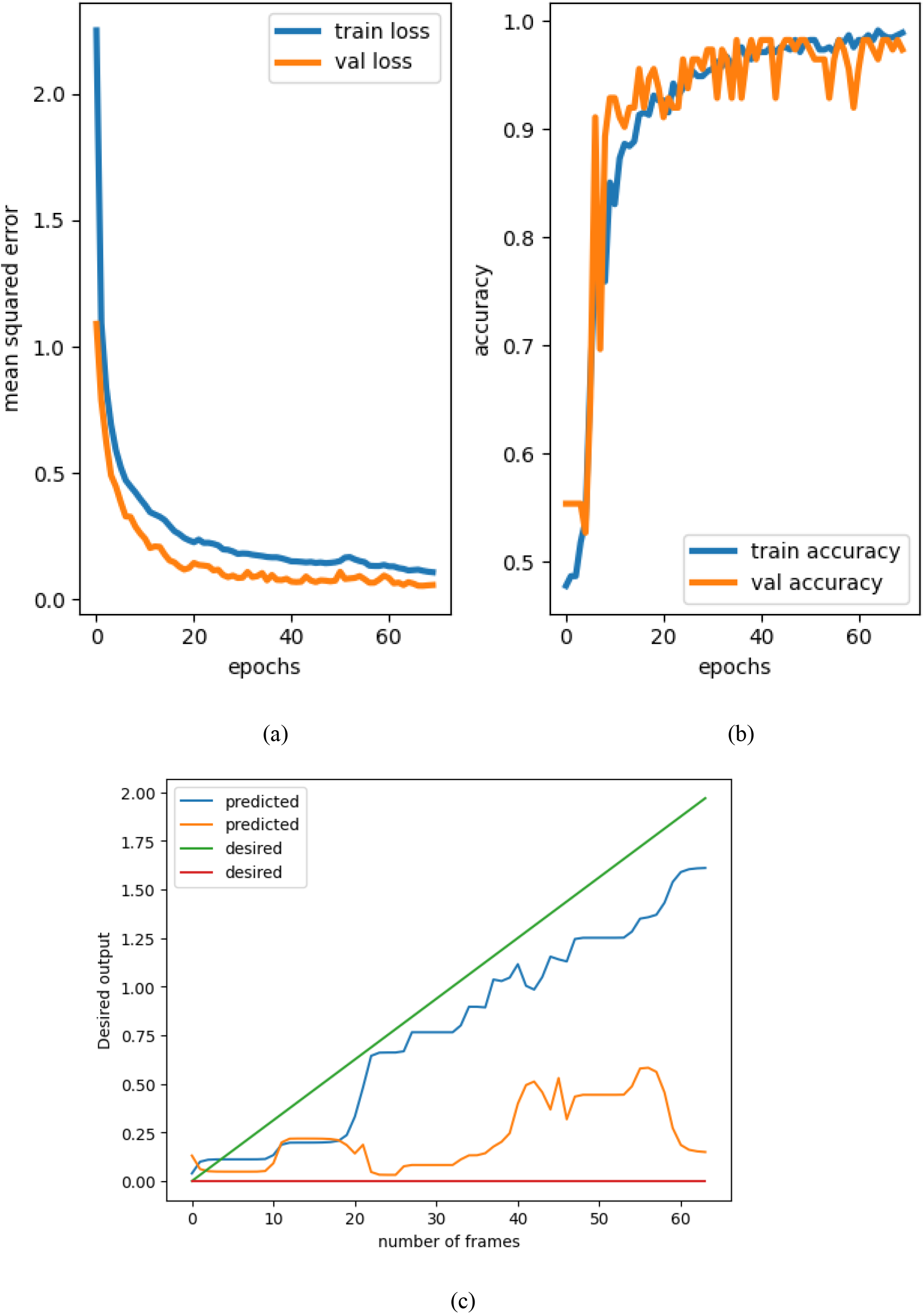
Training -testing performances for sub 15-CHB MIT Dataset (a-b), Plot of output target with desired (c).

**Figure 12:**
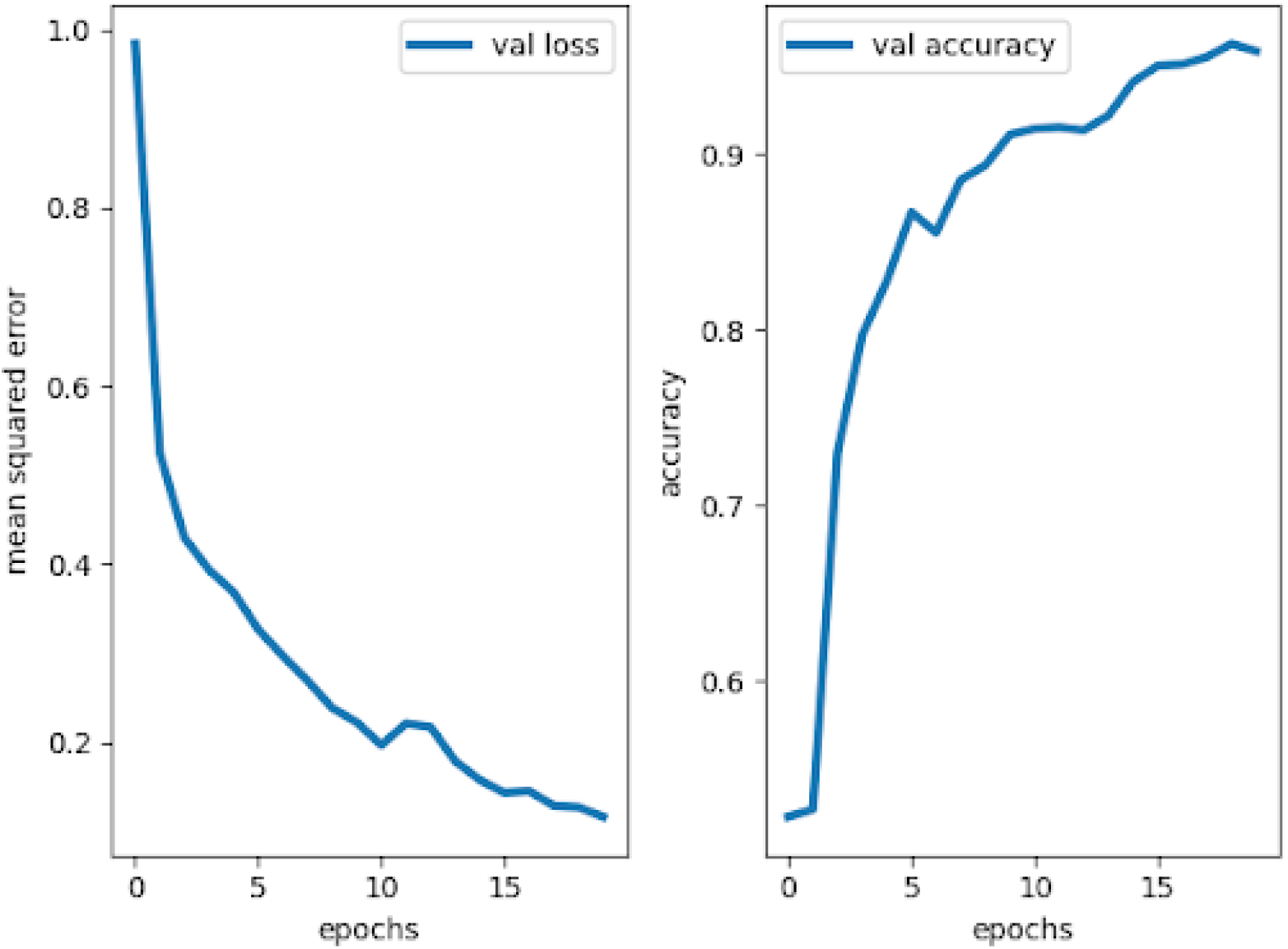
Our proposed model performance on CHB-MIT for the 15th subject during testing (subject-independent case)

#### 4.2.4 Subject-Independent classification on CHB-MIT Dataset using OCNN-Topography

For this task, of all 23 subjects of CHB-MIT dataset, we use 22 subjects for training and hold out of one subject for testing. Model parameters mentioned in Table 8 are used for this task. The average model performance across all patients is reported (Fig 13). We reported a comparison table on previously reported works on subject-specific classification accuracies on the CHB-MIT dataset (Table 10).

**Fig 13:**
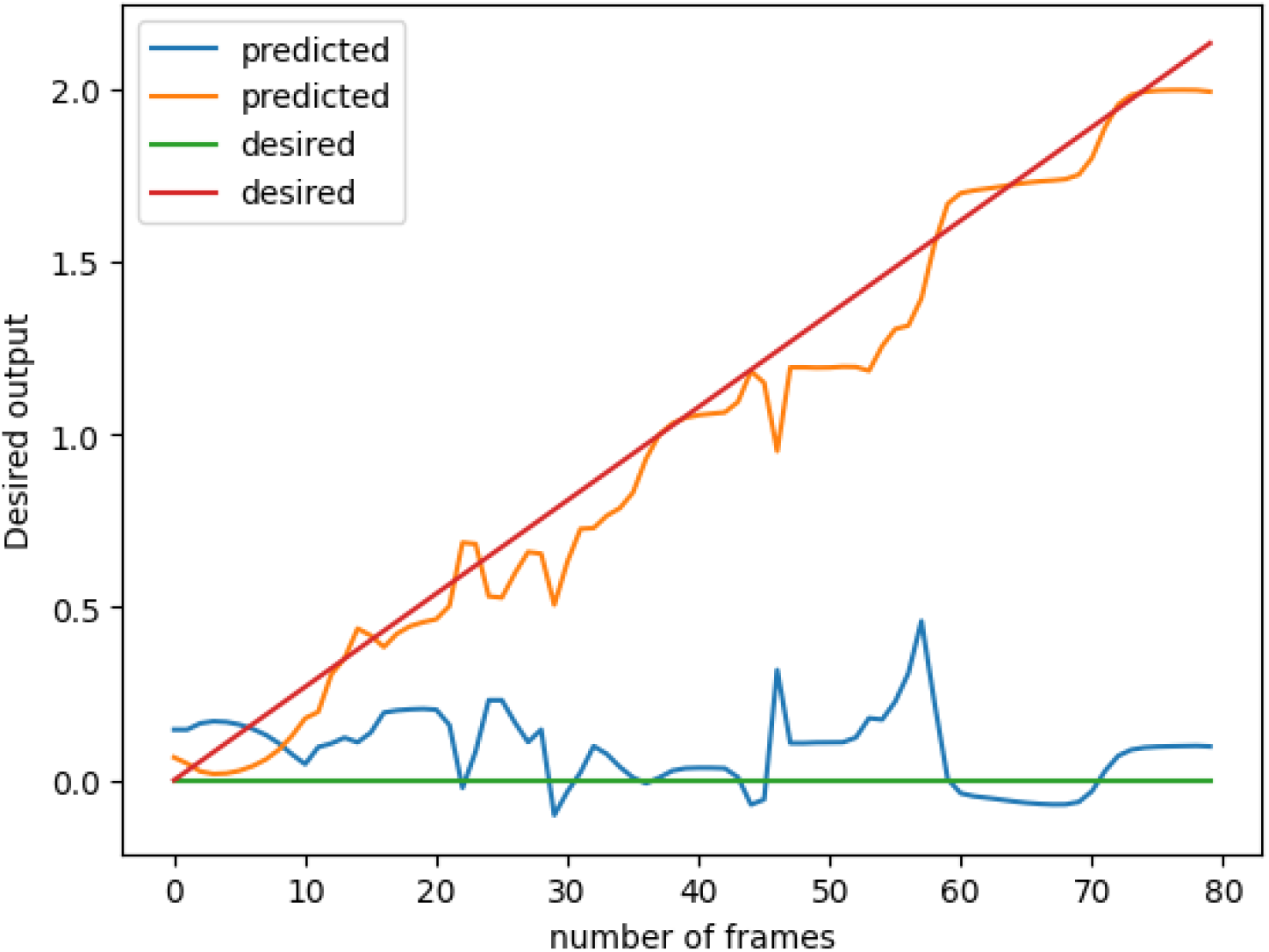
Plot of output target with desired for subject 4 two class motor imagery classification

#### 4.2.5 Motor Imagery Data classifier with OCNN

Classification of human motion intention using the machine learning approach is the most common challenge in Brain-computer interface (BCI) fields. Several feature extractions (wavelet transform [69], Hilbert transform [70], short-time Fourier transform [71]) and channel selection algorithms with recurrent type networks [66-68] have already been applied to decode motor movement from EEG signals.

In this study, each dataset consists of approximately 300 samples of left-hand movement and 300 samples of right-hand movement. Each sample is of 1 sec duration (200 time point). All the 19 EEG channels were used here. We applied 4 samples (20 millisec) non-overlapping window averaging over the data. Then the dataset size becomes 50*19 for each sample. Topoplot is calculated at each time instant. As before, classification is done on an individual subject basis. The topoplot-generated images are first converted from RGB to grayscale and then resized to 50*50 size. The corresponding target output size is also set to (50*2), where the desired output of the correct class increases linearly whereas the other outputs corresponding to the incorrect classes remain zero. The ramp function is generated up to a maximum value of 2. The number of data points present in the ramp function is equal to the number of frames in the video. Therefore, for instance, in general, the input data size per sample is 50*50*50.

Average accuracy over 7 subjects= 81.42%

Compared to the previously reported results on the two-class left vs right-hand movement classification the proposed OCNN network produced an accuracy of 81.42% over 75% reported in [36].

## Discussion

In this study, we present a Deep Oscillatory Neural Network (DONN) that can perform prediction as well as classifications of various classes of EEG signals. We also present a convolutional version of DONN (OCNN) which can classify the sequence of scalp topoplot images using its spatiotemporal filters.

In our previous study, we have shown a purely generative model using an oscillatory network consisting of Hopf oscillators, where we predict and generate EEG time series of various sleep stages [42]. In that model, we have used a layer of oscillatory neurons as a generative layer, followed by a layer of sigmoid neurons that can produce accurate reconstructions of high-dimensional EEG data. Recently we have developed DONN and OCNN models which can model input-output behaviors for both classification and regression using simple data sets.

In the current study, we demonstrate the classification and regression/prediction of EEG signals using DONN. For the regression problem, we demonstrated accurate prediction of the nearest EEG channel’s data with an average RMSE error of 0.01. Two variations of the application have been shown in this study where in the first case, all training and testing are done on an individual subject basis and in the second case, training is done on 4 subjects and testing is done on the 5^th^ subject. There are a large number of studies on EEG channel reduction using data compression methods like PCA (Principal Component Analysis), ICA (Independent Component Analysis) etc. [56-58]. However, most of the algorithms have huge computational overhead and are time-consuming. In this work, we also showed one more possible application of the DONN model viz., the nearest channel prediction method, which could be an effective method for EEG data compression.

Along with prediction, the proposed DONN and OCNN networks can classify different EEG datasets with higher accuracy than previously reported performances in the literature (Table 9, Table 10). Existed methods are quite time-consuming, larger network size and larger trainable parameters. A comparison table has been shown in Table 9-Table 10.

**Table 9:** Performance comparison of the proposed OCNN model with previously published results for the CHB MIT Dataset.

| Classifier Approach | Model performances | Size of Networks/parameters |
| --- | --- | --- |
| [32] a) MLP<br>b) DCNN+MLP<br>c) DCAE+Bi-LSTM<br>d) DCAE+Bi-LSTM+CS | a) 83.63<br>b)94.10%<br>c) 99.66%<br>d) 99.66%<br>e) 99.66% | Number of trainable parameters in each case:<br>a) 8,870,291<br>b)520477<br>c) 27657<br>d) 27657<br>e) 18345 |
| [33] Deep metric learning | 86.68% | 7 convolution layer (4,24,24,72,128,256 map), number of hidden layers 3, neuron: 64,8,1 . |
| [34] ST-MLPs | sensitivity-96.6,<br>specificity-86.7,<br>AUC-93.8%, | Number of trainable parameters-70k |
| [28] Adaptive Haar Wavelet-based Binary Grasshopper Optimization Algorithm and Deep Neural Network (AHW-BGOA-DNN) | 93.24% |  |
| [44] Bidirectional LSTM | 83.89% | Model parameters: 197078 |
| [45] Independent RNN | 87% | 128 neuron -1st 5 RNN layer<br>200 neuron-2nd 5 RNN layer<br>250 neuron-3rd 5 RNN layer<br>Fully connected layer-100 neuron<br>epoch-30,<br>trainable parameter not mentioned |
| Our proposed model | **95.98** | Model parameters:<br>177982 |

**Table 10:**
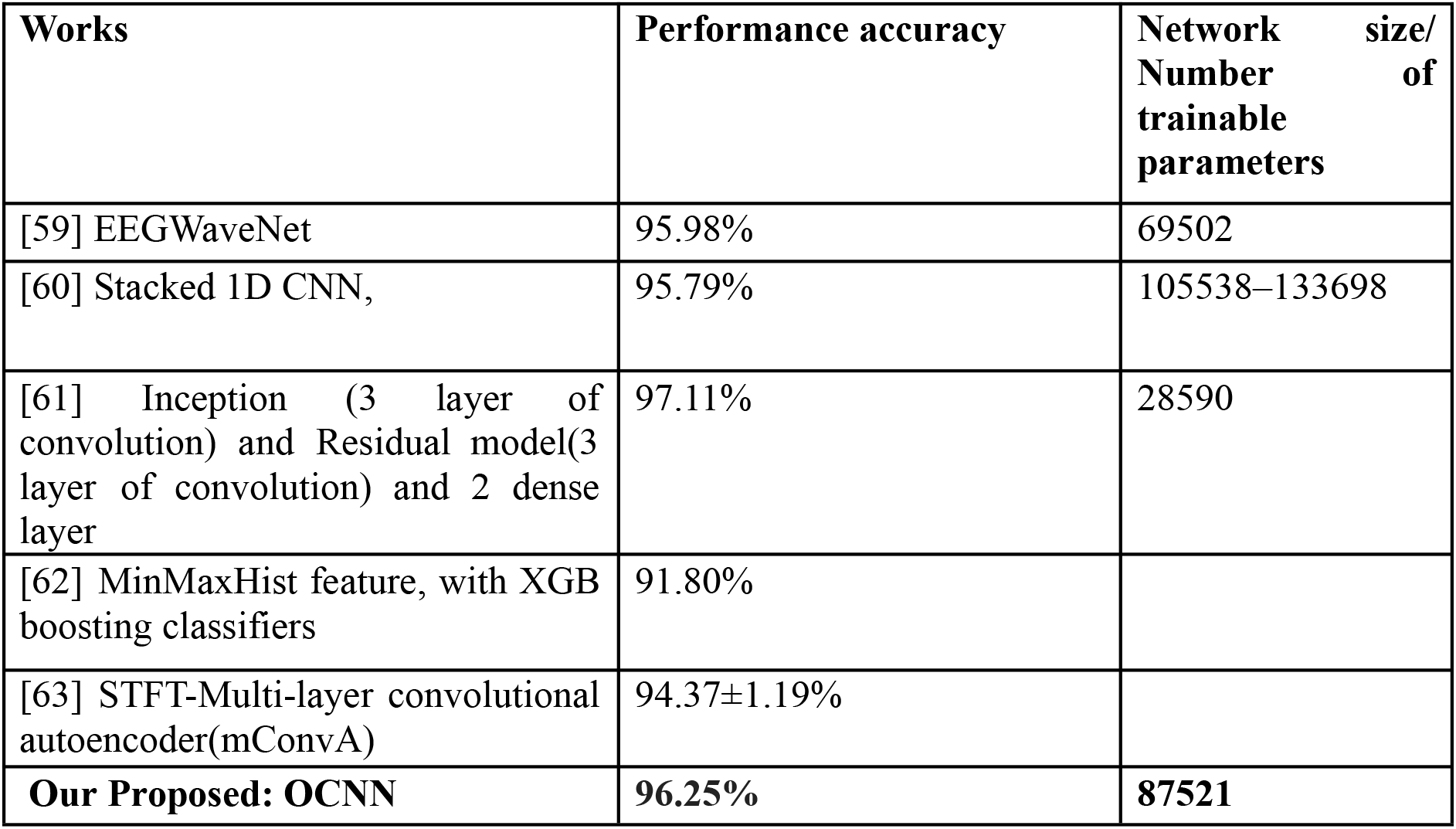
Performance comparison of the proposed OCNN model with previously published results for the CHB MIT dataset (subject independent case).

**Table 10:** Details of the OCNN model for motor imagery data.

| Model parameters |  |
| --- | --- |
| Image size | 50,50 |
| Frames of the video | 50 |
| Number of kernels for F1 layer | 5 |
| kernel dimension of F1 layer | 5,5 |
| After 1<sup>st</sup> convolution size of feature map | 5,46,46 |
| Size of oscillators (OS1) | 5,46,46 |
| Number of kernels for F2 layer | 5 |
| kernel dimension of F2 layer | 5,5 |
| Feature map dimension after F2 layer | 5,42,42 |
| H1 (Dense layer neuron) & o/p layer neuron | 15 & 2 |
| Drop out | 0.05 |
| Learning rate | 0.001 |
| optimizer | adam |

**Table 11:** OCNN model performances for 7 subjects.

| Subject | Accuracy |
| --- | --- |
| Subject 1 | 75% |
| Subject 2 | 86.67% |
| Subject 3 | 85.83 |
| Subject 4 | 90.83% |
| Subject 5 | 73.33% |
| Subject 6 | 78.29% |
| Subject 7 | 80.00 |

In this work, we introduce Topoplot maps as a sequence of images that are classified by our novel OCNN, and as far as of authors’ knowledge there is no other study that uses a sequence of topographic images for EEG classification. Here, we have shown two examples: a) CHB-MIT, and b) Motor imagery dataset using spatiotemporal topographic video and OCNN classifier. Previously topographic images have been classified by CNN type networks [64-65], whereas in the present study, we present topographic images as a sequence and detect the seizure event. In [64], the authors used 2D AlexNet CNN which consists of five layers of convolution, four deep layers, and one output layer (sizes of the feature maps in five layers are: 96,256, 364, 364, and 256 and the kernel sizes in each layer are: 11*11, 5*5, 3*3, 3*3, and 3*3 & the sizes of the four deep layers are: 1000,500,250,125 and 4 neurons). Compared to that model [64], our network can perform better with lesser numbers of network parameters. In addition, OCNN model helps to extract temporal and spatial features from topograph sequences by which gives more accurate classification accuracy. Also, we perform both patient-specific and patient-independent seizure detection on the CHB-MIT dataset using OCNN. Both have their unique advantages: patient-specific study is useful for event detection (occurrence of seizure events) whereas patient-independent analysis is more useful for diagnosis. The method described in this study can be various healthcare problems (i.g: epilepsy patients, stroke patients, mental disorders and computer brain-computer interfaces etc.).

In future works, for more complex, sequential classification problems (like hybrid behavioral and classifier model, multi-class classification problems etc.), which involve decision-making over very long sequences, we can add memory elements like flip-flop neurons [76] to our DONN or OCNN models. The study shows the potential application of deep oscillatory neural networks and learning algorithms for classification raw EEG data without feature extraction, which introduces a new perspective to facilitate clinical decision-making.

## Acknowledgement

We acknowledge the financial support from DBT Project, Govt. of India, and Center for Complex Systems and Dynamics, IIT Madras. The authors acknowledge the support of fellow lab mates Sheth Jenil Samir, and Priyavrata Tiwari for preprocessing the EEG signals.

