## Supplementary material for "Electroencephalogram (EEG) Classification using a bio-inspired Deep Oscillatory Neural Network": supplimentary

### Sleep Data classification using DONN Network:

A. Physionet Sleep Dataset[5]

This database consists of whole-night polysomnogram recordings of 197 subjects [16]. Polysomnogram recording contains EEG, EMG, and chin EMG. EEG datasets are recorded from two electrodes (Fpz-Cz and Pz-Oz). The Datasets are in 'edf' format and corresponding sleep stage scoring files are also attached with the main datasets. This dataset is band pass filtered between (0.1 to 20 Hz).

We created 10s batch of EEG Fpz-Cz for sleep stages N3 and REM from psg data using duration from hypnogram data for only those subjects who had at least 300 batches and concatenate the 300 batches from both stages together followed by random shuffling to avoid any bias in our analysis.
We performed a 0.8 train-test split on our data of length 600. On training our model with 80 relu neurons and 80 oscillator neurons (hopf oscillator), then 40 and 30 tanh neuron layers successively, we obtained a training accuracy of 97.58% and a test accuracy of 96.67% average of 5 subjects.

| | |
|---|---|
| [2] 1D-CNN | 97.85 |
| [3] VG-HVG+SVM | 97.9 |
| [4] Wavelet filter + SVM | 97.8 |
| **Our proposed** | **96.67** |

Table 4: Comparison of accuracies with existing literature compare to our propose DONN network for physionet epilepsy Dataset.

Claustrophobic EEG Data classification using OCNN [1]

EEG data has been collected from 9 claustrophobic volunteers and 13 healthy volunteers using the Mitsar 31-channel EEG system [1]. All the EEG data was collected in an open rest state condition. In one case EEG data was collected in a large-spaced laboratory. In 2nd case, data was recorded from a medium-spaced room. In 3rd case, EEG data acquisition was performed in the same chamber but with reduced dimensions.

In this study we have taken 5 claustrophobic subjects (S01, S03, S12, S13, S14) EEG as well as five 5 healthy subjects (S02,S04,S05, S06, S07) EEG for all three experimental condition. The dataset is recorded with 2000 Hz, further, it is down-sampled to 500 Hz. We applied bandpass filtering from the total 300-sec data (0.5 to 40 Hz) and extracted 100th to 200th sec (100 sec EEG-50000 sample points) from each dataset. The rest of the part of data we ignored in this study. Then a 50-sample window (0.1 sec) is applied to the dataset. So, it creates 1000 chunks of data each having a length of 50 samples and 31 channels.

| Recording condition | Testing accuracy |
|---|---|
| R0 | 95.25 |
| T1 | 91.17% |
| T2 | 92.17% |

Supplimentary References:

[1] Moradi, D., Eyvazpour, R., Rahimi, F., Jahan, A., Rasta, S. H., & Esmaeili, M. (2021). Electroencephalographic activity in patients with claustrophobia: a pilot study. *Journal of Medical Signals & Sensors*, *11*(4), 262-268.

**[2]** Yildirim, O., Baloglu, U. B., & Acharya, U. R. (2019). A deep learning model for automated sleep stages classification using PSG signals. *International journal of environmental research and public health*, *16*(4), 599.

**[3]** Zhu, G., Li, Y., & Wen, P. (2014). Analysis and classification of sleep stages based on difference visibility graphs from a single-channel EEG signal. *IEEE journal of biomedical and health informatics*, *18*(6), 1813-1821.

[4] Sharma, M., Goyal, D., Achuth, P. V., & Acharya, U. R. (2018). An accurate sleep stages classification system using a new class of optimally time-frequency localized three-band wavelet filter bank. *Computers in biology and medicine*, *98*, 58-75.

[5] Kemp, B., Zwinderman, A. H., Tuk, B., Kamphuisen, H. A., & Oberye, J. J. (2000). Analysis of a sleep-dependent neuronal feedback loop: the slow-wave microcontinuity of the EEG. *IEEE Transactions on Biomedical Engineering*, *47*(9), 1185-1194.
